# Introgressive hybridization as a potential source of evolutionary rescue in an extreme peripheral environment

**DOI:** 10.64898/2026.07.24.740513

**Authors:** Sahar Javaheri Tehrani, Yu-Chi Chen, Kees van Oers, Mirte Bosse, Martin Päckert, Jochen Martens, Mansour Aliabadian, Niloofar Alaei Kakhki, Toni I. Gossmann

## Abstract

Range-margin populations often experience demographic and environmental conditions that constrain their adaptive potential. Genetic drift can erode diversity, while gene flow from the range core may swamp locally favoured alleles. These limitations may be severe in extreme environments, yet how peripheral populations persist and adapt there remains poorly understood. Introgression from neighbouring lineages may address both constraints by adding genetic variation and introducing alleles pre-tested by selection, potentially contributing to evolutionary rescue. Here, we analyse 112 whole genomes across the great tit (*Parus major* ) complex to investigate how peripheral populations persist and adapt, focusing on *P. m. intermedius*, a grey-plumaged population in arid northeastern Iran at the southeastern margin of the yellow-breasted *major* lineage, bordering the grey *bokharensis* lineage. We found that *intermedius* falls within *major* in the first nuclear species tree of the complex and the mitochondrial phylogeny. As expected for a peripheral population, it is genetically differentiated, yet retains relatively high nucleotide diversity and carries low-level genomewide *bokharensis* ancestry. Against low genome-wide differentiation from *major*, a 35-kb region on chromosome 24 shows excess allele sharing with *bokharensis* up to six-fold above the genome-wide average, a *bokharensis* origin in local genealogies and ancestry inference, and reduced diversity and negative Tajima’s *D* consistent with positive selection. This region contains *BCO2*, a carotenoid-cleaving gene, with differentiated 5*^′^* untranslated-region variants, implicating regulatory variation in carotenoid-based plumage. Our findings suggest that introgression from closely related lineages can buffer peripheral populations against loss of genetic variation and facilitate adaptation to harsh, divergent habitats beyond the range core, highlighting hybridization as an underappreciated source of resilience at species’ range limits.

**Significance Statement:** Adaptation often fails at range margins, where populations are expected to be differentiated yet genetically depleted, and gene flow from the core can swamp locally favoured alleles. Introgression could counter both, buffering by adding variation and enabling adaptation through pre-tested alleles. We examined these roles in *Parus major intermedius*, a peripheral great tit population in the putative contact zone between the yellow Eurasian *major* and grey *bokharensis* lineages. We analysed 112 genomes from across Eurasia; the first nuclear species tree places *intermedius* within *major*. *Intermedius* is differentiated yet retains most of its diversity and carries low-level *bokharensis* ancestry, while one region stands out: containing the carotenoid-cleaving gene *BCO2*, it retains *bokharensis* ancestry far above genome-wide levels, consistent with selection. More broadly, hybridization with neighbouring lineages may be an overlooked source of variation and adaptive potential for range-margin populations facing extreme or changing environments.

## Introduction

Why species cannot keep adapting along environmental gradients, and thus expand their ranges indefinitely, remains a central question in evolutionary biology (Kirkpatrick and Barton 1997; Bridle and Hoffmann 2022). At the heart of the problem lies gene flow, which pulls in opposite directions: it supplies the genetic variation on which adaptation depends, yet it can also swamp local adaptation with alleles suited to other environments (Lenormand 2002). Haldane first suggested that such swamping, by reducing fitness and hence population size at the edge, could set a sharp limit to a species’ range (Haldane 1956). Theory now suggests that the decisive factor is genetic variance itself: when variance can evolve, gene flow alone need not halt adaptation, but genetic drift in small marginal populations can erode variance below the level required to adapt to spatially varying conditions, producing an abrupt range margin even where the environment changes only gradually (Polechová and Barton 2015). Peripheral populations, which often live under marginal or extreme conditions at the boundary between persistence and extinction (Sexton et al. 2009; Bridle and Hoffmann 2022), therefore face a double bind: depending on their connectivity, they receive either too little gene flow to maintain the variation needed to adapt, or too much to retain locally favoured alleles (Bridle and Vines 2007; Eckert et al. 2008). How, then, can they acquire the variation they need to adapt?

Interspecific introgression, whereby hybridization followed by repeated backcrossing transfers genetic material between species, offers one answer (Edelman and Mallet 2021). Drawing on a far more divergent gene pool than gene flow among conspecific populations, it introduces much greater genetic variance. When the donor lineage is already adapted to conditions beyond the recipient’s range edge, introgression delivers not neutral novelty but variation pre-tested by selection: alleles that may be favoured at the margin rather than swamped, that bypass the waiting time for new beneficial mutations, and that enter the population at higher frequencies than new mutations, reducing their risk of loss by drift (Hedrick 2013; Gaczorek et al. 2024).

Genomic studies have confirmed that introgressed alleles can underlie rapid adaptation, from high-altitude tolerance in Tibetans and seasonal camouflage in snowshoe hares to carotenoid plumage in warblers, and can enable persistence in extreme or newly colonized environments (Huerta-Śanchez et al. 2014; Jones et al. 2018; Baiz et al. 2021; Oziolor et al. 2019; Rendón-Anaya et al. 2021; Jorquera et al. 2025). Such adaptive introgression recurrently involves genes from particular functional categories, notably those underlying immunity (Enard and Petrov 2018), pigmentation (Giska et al. 2019) and tolerance of environmental (Whitney et al. 2006) and anthropogenic stressors (Song et al. 2011), possibly because related species often face similar selective pressures (Gaczorek et al. 2024). Whether introgression can likewise buffer peripheral populations against both drift and swamping, and potentially contribute to their evolutionary rescue (Bell 2017; Brauer et al. 2023), nevertheless remains an open question.

The great tit (*Parus major* ) complex offers an excellent opportunity to address this question. One of the most intensively studied wild vertebrates, it combines decades of ecological data with rich genomic resources (van Oers et al. 2014; Laine et al. 2016; Spurgin et al. 2024), and its European populations have long served as a model for the interplay between gene flow and local adaptation (Postma and van Noordwijk 2005; Perrier et al. 2018; Salmón et al. 2021). Its main lineages (*major*, *minor*, *cinereus*, *bokharensis* and an eastern Himalayan lineage) diversified during the Pleistocene and now meet in well-defined contact zones (Kvist et al. 2007; Song et al. 2020) (Fig. 1A). At the southeastern margin of the *major* lineage, in northeastern Iran, the forested north and west are inhabited by the yellow-breasted *P. m. karelini* and *P. m. blanfordi*, whereas further east, where forests give way to open, arid landscapes, the grey-plumaged *P. m. intermedius* occurs (Fig. 1B; Fig. S1). Because *intermedius* occurs near the presumed contact zone of the *major*, *bokharensis* and *cinereus* lineages and shares the grey plumage of the latter two, it has historically been considered a possible hybrid between the *major* lineage and either *bokharensis* or the *cinereus* group (Vaurie 1950; Formozov et al. 1993; Martens 1996; Eck and Martens 2006). Previous studies based on mitochondrial and microsatellite markers, however, place *intermedius* within the *major* lineage, with only a single individual carrying a *bokharensis* mitochondrial haplotype (Kvist et al. 2007; Javaheri Tehrani et al. 2021). This raises the question of whether the plumage and habitat similarities between *intermedius* and *bokharensis* reflect genome-wide shared ancestry, or whether introgression has introduced genetic variation that potentially contributes to their resemblance. More broadly, this system offers an opportunity to investigate the role of introgression in shaping the genetic composition and evolutionary divergence of peripheral populations at species’ range margins, and in facilitating their rapid adaptation to harsh, divergent habitats beyond the range core.

**Figure 1:**
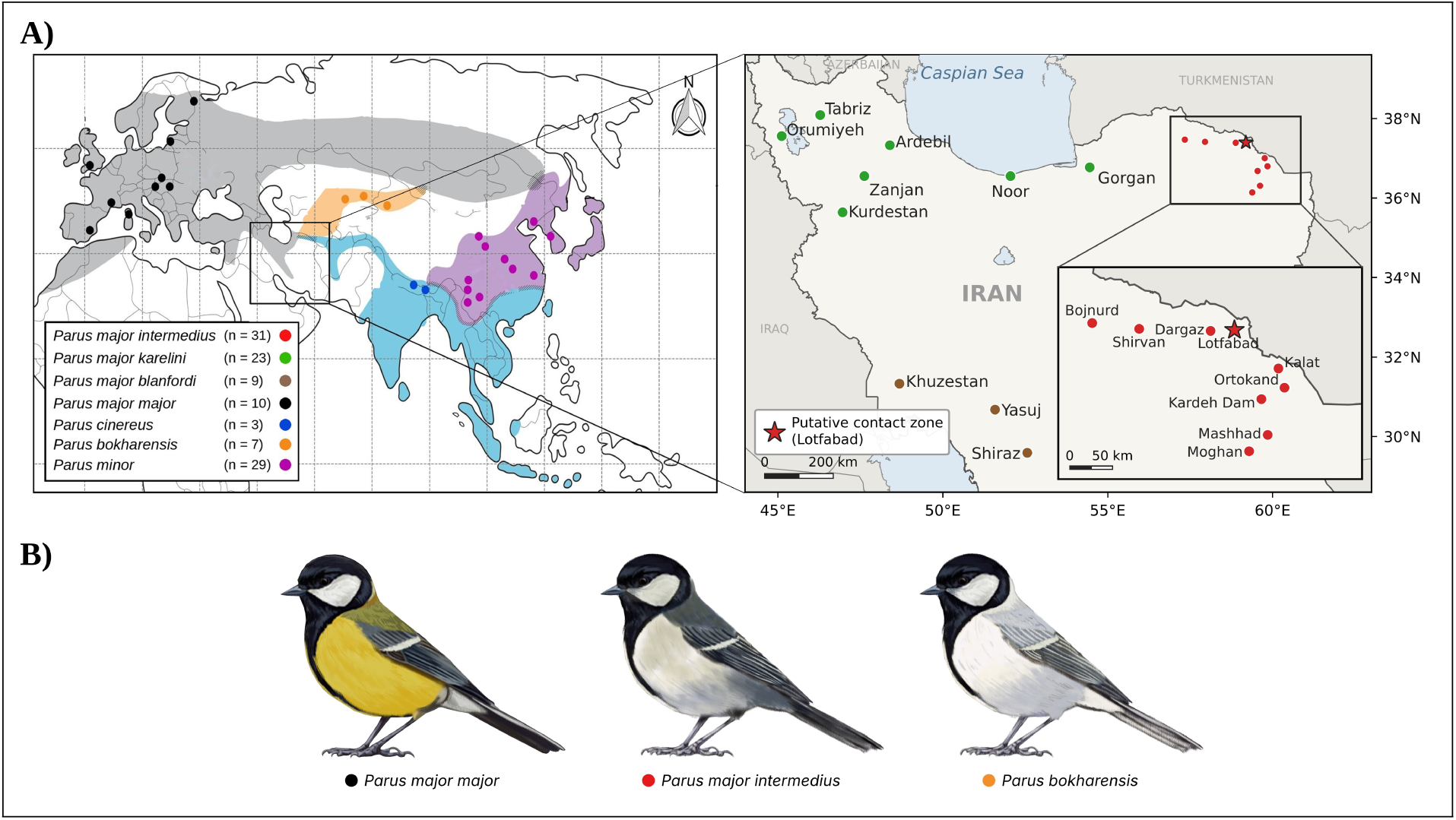
Geographic distribution, sampling design and plumage of the great tit (*Parus major* ) complex. (**A**) Left: approximate distribution ranges of the main lineages across Eurasia, adapted from Martens (1993); hatched areas indicate contact zones between lineages. Sampling localities represent 112 individuals, including 73 newly sequenced and 39 previously published genomes (*∼*3,000 Gb of genomic sequence data). Right: sampling localities in Iran for the three focal subspecies, *P. m. karelini* (green), *P. m. blanfordi* (brown) and *P. m. intermedius* (red); the inset enlarges the *intermedius* range in northeastern Iran. The star marks Lotfabad, the putative contact zone between *intermedius* and *bokharensis*. (**B**) Plumage of *P. m. major*, *P. m. intermedius* and *P. bokharensis*. *Alt text:* Maps of Eurasia and Iran showing the distribution ranges of the great tit lineages, contact zones and sampling localities, including Lotfabad in northeastern Iran, with illustrations of the yellow-breasted *major*, grey *intermedius* and grey *bokharensis*.

Here, we use whole-genome resequencing data from 112 individuals of four lineages of the great tit complex, sampled across their distribution ranges, to (i) reconstruct the first genome-scale species tree of the complex and establish the evolutionary origin of *intermedius*, (ii) quantify gene flow between *bokharensis* and *intermedius*, and (iii) identify alleles preferentially retained from *bokharensis* in *intermedius* and test whether they could account for the morphological similarity between the two. To this end, we begin by resolving relationships within the complex by integrating nuclear and mitochondrial phylogenies with demographic inference. We then trace the extent and genomic distribution of gene flow between *bokharensis* and *intermedius*, combining admixture analyses and tests of excess allele sharing with genome-wide scans of differentiation and introgression. Finally, we pinpoint candidate genes within introgressed regions and dissect their coding and regulatory variation.

## Results

### Discordant mitonuclear phylogenies and genome-wide excess allele sharing

The mitochondrial haplotype network resolved four major haplogroups corresponding to the *major*, *bokharensis*, *minor* and *cinereus* lineages (Fig. 2A). Mitogenome-based phylogenetic analyses recovered the same four lineages as well-supported monophyletic clades (Fig. 2B; Fig. S2). The mitochondrial phylogeny placed *major* and *bokharensis* as sister lineages (bootstrap support 97), together forming a clade that was sister to a second clade comprising *minor* and *cinereus* (98). Within the mitochondrial tree, *intermedius*, together with a subset of *karelini* individuals, formed a distinct mitochondrial subclade that was clearly separated from the main *major* clade with strong bootstrap support (Fig. S2). Notably, two *intermedius* individuals from the putative contact zone (Lotfabad, Iran–Turkmenistan border) between the *major* and *bokharensis* lineages carried *bokharensis* mitochondrial haplotypes (Fig. 2A). Conversely, two museum specimens assigned to *bokharensis*from Kazakhstan carried *major* -associated mitochondrial haplotypes (Fig. S2).

**Figure 2:**
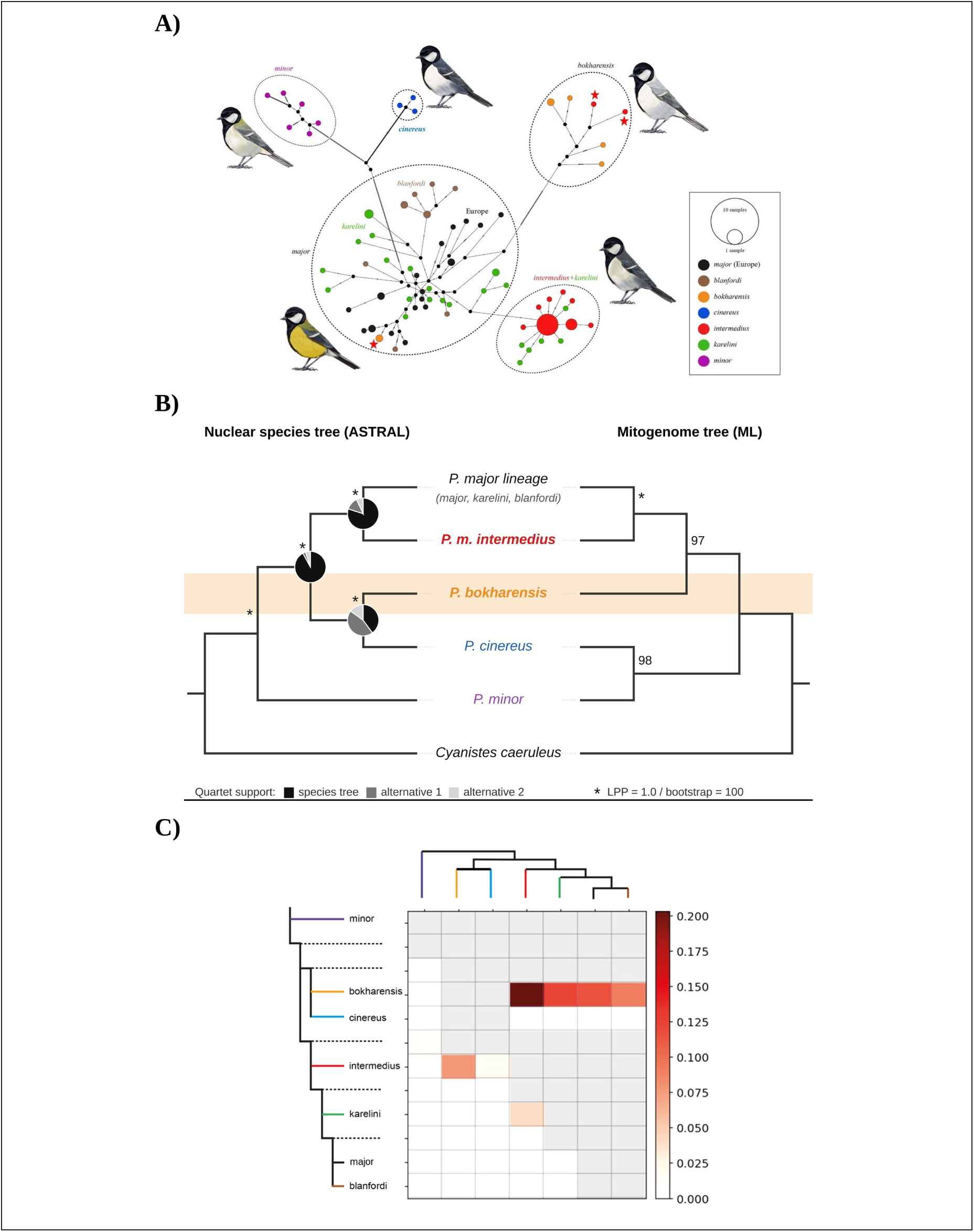
Mitonuclear discordance and excess allele sharing in the *Parus major* complex. **(A)** TCS network of whole mitogenomes; circle size reflects haplotype frequency, and stars mark individuals with discordant haplotypes. **(B)** ASTRAL-III nuclear species tree (left) and schematic mitogenome tree (right; full tree in Fig. S2), with *major*, *karelini* and *blanfordi* combined as the *major* lineage and *Cyanistes caeruleus* as outgroup. Pies show quartet support on internal branches (q1, black; q2, dark grey; q3, light grey); asterisks, LPP = 1.0 or bootstrap = 100; numbers, bootstrap values. The orange band highlights the discordant placement of *bokharensis*. **(C)** *f* -branch statistics (*f_b_*; Dsuite) between branches (y-axis) and taxa (x-axis); grey cells, tests not possible under the topology. *Alt text:* A mitochondrial haplotype network of the great tit complex; a nuclear species tree compared with a mitogenome tree, showing *bokharensis* as sister to *cinereus* in the nuclear tree but to *major* in the mitochondrial tree; and a heatmap of *f* -branch excess allele sharing, highest between *bokharensis* and the *major* lineage, especially *intermedius*.

The species tree inferred from genome-wide nuclear gene trees was largely concordant with the mitochondrial phylogeny but differed in the placement of *bokharensis* (Fig. 2B). In the nuclear species tree, *bokharensis* was sister to *cinereus*, whereas in the mitochondrial tree it was sister to *major*. All relationships in the nuclear species tree received maximal support (local posterior probability = 1.0; Fig. 2B). Quartet support was high for the internal branch separating the *major* lineage, including *intermedius*, from *bokharensis* and *cinereus* (q1 = 0.87), and for the placement of *intermedius* as sister to the remaining *major* lineage (q1 = 0.69). In contrast, the branch uniting *bokharensis* and *cinereus* showed substantial gene tree discordance: only 43% of quartets supported this relationship, and the two alternative topologies were unequally represented (q2 = 0.36, q3 = 0.21).

Genome-wide introgression tests revealed heterogeneous excess allele sharing across the complex (Fig. 2C). The strongest *f* -branch signals involved the *bokharensis* branch, which showed excess allele sharing with all lineages of the *major* clade. Values were highest with *intermedius* (*f_b_ ≈* 0.20), intermediate with *karelini* and *major* (*f_b_ ≈* 0.14) and lowest with *blanfordi* (*f_b_ ≈* 0.10).

### Contrasting demographic histories of the *major* and *bokharensis* lineages

Historical changes in effective population size (*N_e_*) inferred with MSMC2 revealed contrasting demographic trajectories for the *major* and *bokharensis* lineages (Fig. S3). After a gradual decline between *∼*2 and *∼*1 Mya, the effective population size of *major* increased approximately twofold, from *∼*3 *×* 10^5^ at *∼*1 Mya to a peak of *∼*7 *×* 10^5^ at *∼*200 kya, followed by a moderate decline towards the most recent time intervals. In contrast, *bokharensis* declined steadily from *∼*5 *×* 10^5^ at *∼*2 Mya and has remained small and stable at *∼*1 *×* 10^5^ over the last *∼*300 kyr.

### *Intermedius* genomes reflect asymmetric admixture with *bokharensis*

In the PCA including all taxa, PC1 (27.3%) separated *minor* from all other lineages, and *intermedius* clustered with the *major* lineage (*major*, *karelini* and *blanfordi* ; Fig. S4A). When *minor* was excluded, PC1 (16.8%) separated the *major* lineage from *bokharensis* and PC2 (9.4%) separated *cinereus* (Fig. S4B). Along PC1, nine of eleven individuals from Lotfabad were displaced towards *bokharensis*, whereas individuals from all other localities clustered with the *major* lineage; no *intermedius* individual was displaced towards *cinereus*.

A regional PCA of the three focal lineages showed the same pattern (Fig. 3A). PC1 (20.29%) separated *major* from *bokharensis*, with most *intermedius* individuals clustering close to *major* ; the *intermedius* individuals from Lotfabad, in the putative contact zone, were shifted towards *bokharensis* along this axis. PC2 (4.92%) distinguished *intermedius* from *major*, indicating that *intermedius* forms a distinct genetic cluster rather than a continuum between *major* and *bokharensis*.

**Figure 3:**
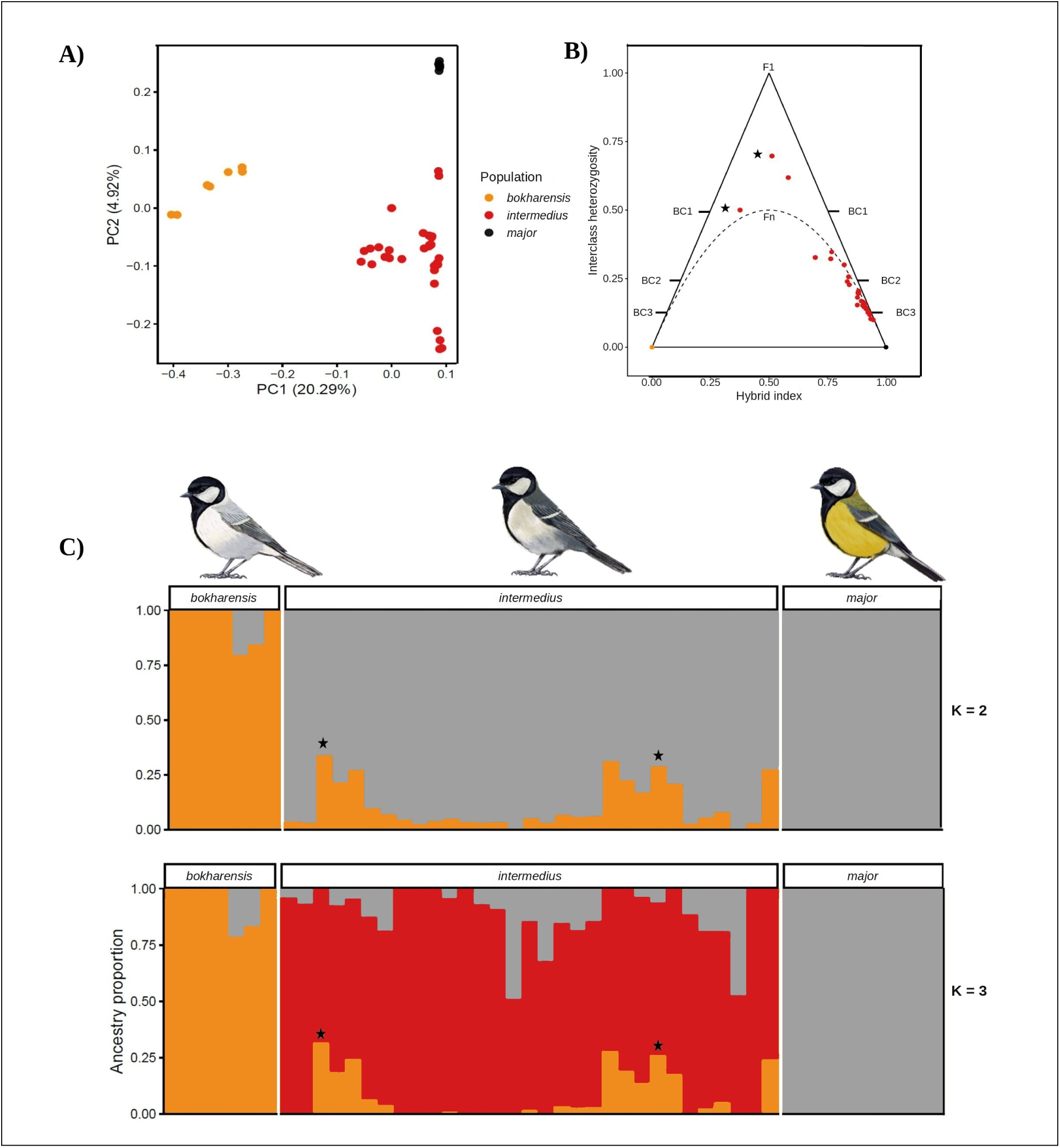
Genome-wide patterns of admixture between the *major* and *bokharensis* lineages. **(A)** Principal component analysis (PCA) of *major*, *intermedius* and *bokharensis* individuals (regional data set; see Fig. S4 for all lineages). **(B)** Hybrid index versus interclass heterozygosity. **(C)** ADMIXTURE ancestry proportions at *K* = 2 and *K* = 3 (orange, *bokharensis*; grey, *major* ; light red, *intermedius*-specific component). Stars mark the two Lotfabad individuals carrying *bokharensis* mitochondrial haplotypes (B, C). See Fig. S5 for *K* = 2–4 with sample labels. *Alt text:* A PCA, a hybrid index triangle plot and ADMIXTURE bar plots showing that *intermedius* clusters near *major* with small proportions of *bokharensis* ancestry, highest in individuals from Lotfabad.

Hybrid index–heterozygosity analyses further supported admixture in *intermedius* individuals (Fig. 3B). Most *intermedius* individuals exhibited hybrid index values skewed toward the *major* lineage combined with low interspecific heterozygosity, consistent with later-generation hybrids and backcrossed individuals. Only three *intermedius* individuals showed higher interspecific heterozygosity (*≥*0.50) together with intermediate hybrid index values, suggesting early-generation hybrids. Two of these individuals were sampled at Lotfabad and were the same individuals that carried *bokharensis* mitochondrial haplotypes (Fig. 2A).

ADMIXTURE analyses supported the patterns observed in the PCA (Fig. 3C). At *K* = 2, ancestry proportions reflected the primary genomic split between *bokharensis* and the *major* lineage. *Intermedius* individuals exhibited variable proportions of *major* and *bokharensis* ancestry, which were spatially structured across the sampling region: individuals sampled closer to the distribution of *bokharensis* showed higher *bokharensis* ancestry, whereas those located nearer the *major* range showed increased *major* ancestry. The two Lotfabad individuals identified as early-generation hybrids showed the highest *bokharensis* ancestry. At *K* = 3, a third ancestry component emerged that was predominant in *intermedius* individuals, distinguishing *intermedius* from both *major* and *bokharensis*.

### Low genome-wide differentiation is interrupted by localized outliers

Genome-wide differentiation between *intermedius* and European *major* was moderate (*F*_ST_ *≈* 0.056). To identify genomic regions deviating from this genome-wide level, we examined differentiation in sliding windows along the genome. Genome-wide scans of sliding-window *F*_ST_ between *major* and *intermedius* revealed generally low background differentiation punctuated by discrete genomic outlier regions (Fig. 4A). Elevated differentiation was largely restricted to two chromosomes, 6 and 24, with chromosome 24 exhibiting one of the strongest and most localized signals. We therefore focused subsequent fine-scale analyses on chromosome 24 (NC 031792.1). Fine-scale analyses identified a localized region of elevated window-based differentiation around 1.5 Mb. Although most windows across the chromosome showed low differentiation, a narrow peak within this interval exceeded the genome-wide top 1% *F*_ST_ threshold (*F*_ST_ = 0.160), with local peak values of about 0.65 (Fig. 4B). To further resolve local differentiation, SNP-level *F*_ST_ analyses were conducted across the chromosome 24 candidate region. Among 3,381 analyzed SNPs, most sites showed low to moderate differentiation, whereas highly differentiated SNPs (*F*_ST_ *>* 0.9) were concentrated within a 34.9 kb interval (1,490,771–1,525,695 bp; Fig. 4C), hereafter referred to as the candidate interval. Absolute sequence divergence () between *major* and *intermedius* was moderately elevated across the same interval relative to the chromosome-wide average, although comparable values also occurred elsewhere on the chromosome (Fig. 4B).

**Figure 4:**
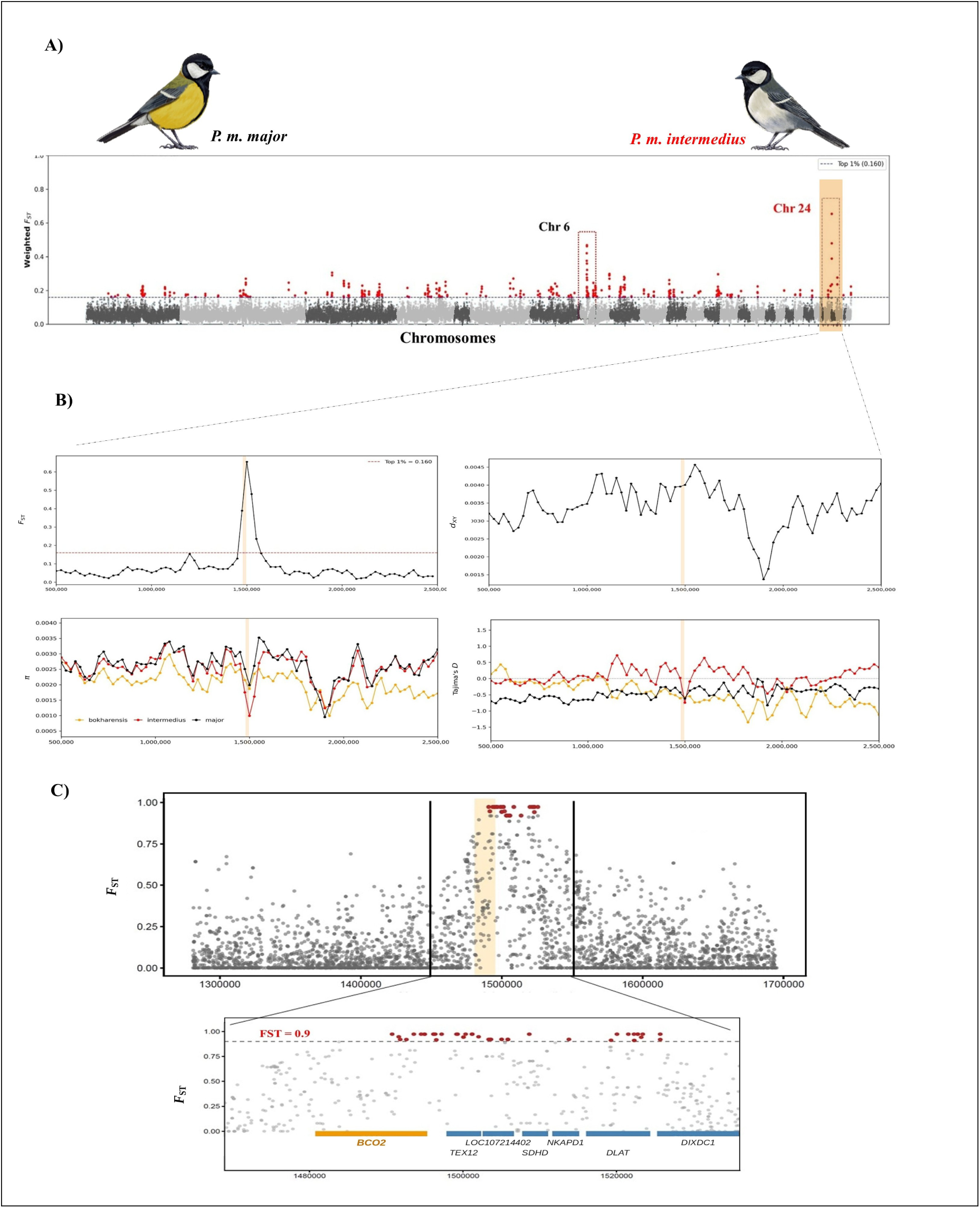
Genome-wide differentiation and fine-scale analysis of a candidate region on chromosome 24. **(A)** Genome-wide distribution of *F*_ST_ between *P. m. major* and *P. m. intermedius*. Points represent sliding windows (50 kb window size, 25 kb step size) across the genome, with outlier windows highlighted in red; the dashed line indicates the top 1% empirical threshold. **(B)** Fine-scale patterns across chromosome 24. Sliding-window estimates (50 kb windows, 25 kb step size) of *F*_ST_, *d_XY_* , nucleotide diversity (*π*), and Tajima’s *D*; *π* and Tajima’s *D* are shown for *major* (black), *intermedius* (red), and *bokharensis* (orange). **(C)** SNP-level *F*_ST_ around the outlier region (top) and across the candidate interval (bottom); SNPs with *F*_ST_ *>* 0.9 are shown in red, with annotated protein-coding genes below. The orange band marks the candidate interval containing *BCO2*. *Alt text:* A genome-wide scan of *F*_ST_ between *major* and *intermedius* with a sharp peak on chromosome 24, and fine-scale plots of *F*_ST_, *d_XY_* , nucleotide diversity and Tajima’s *D* around the candidate interval containing *BCO2*.

Genome-wide nucleotide diversity (*π*) was broadly similar between *major* and *intermedius*, whereas *bokharensis* exhibited consistently lower diversity across the genome (Fig. S6B). Within the chromosome 24 outlier region, *π* was locally reduced in all three focal lineages relative to flanking regions, with the strongest reduction observed in *intermedius* (Fig. 4B).

Genome-wide Tajima’s *D* showed contrasting lineage-specific patterns, with predominantly positive values in *intermedius* and negative values in both *major* and *bokharensis* (Fig. S6A). Within the chromosome 24 outlier region, Tajima’s *D* in *intermedius* shifted toward negative values (about *−*0.7) relative to surrounding regions, whereas *major* remained consistently negative and *bokharensis* showed increasingly negative values across the same region (Fig. 4B).

### A localized ancestry shift links *intermedius* to *bokharensis*

To test whether the differentiated region on chromosome 24 in *intermedius* reflects introgression from *bokharensis*, we applied three complementary approaches: topology weighting, *f_dM_* , and genome polarization (Fig. 5; see also Fig. S7 for genome-wide *intermedius*–*bokharensis* differentiation).

**Figure 5:**
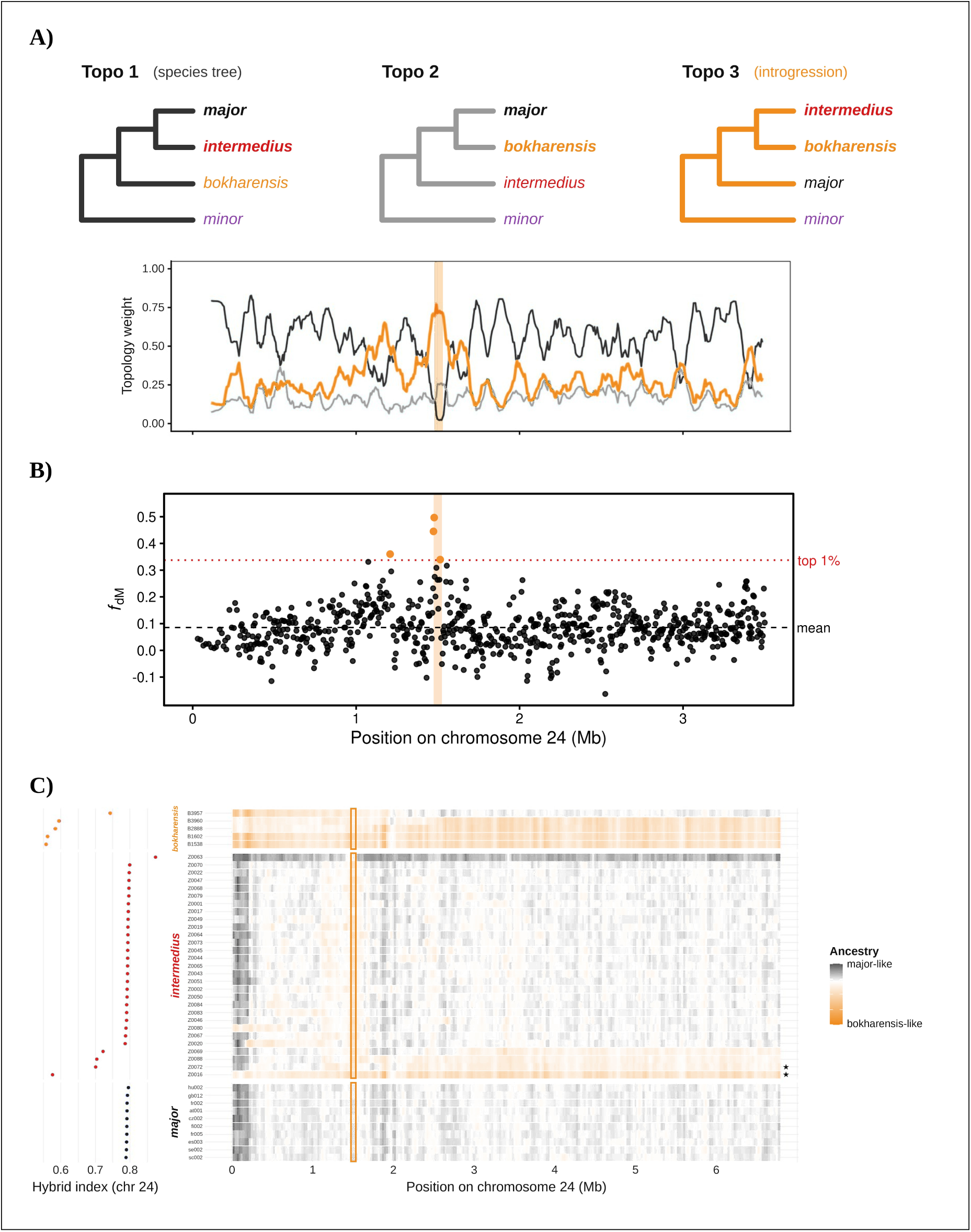
Evidence of introgression from *bokharensis* into *intermedius* on chromosome 24. **(A)** Topology weighting (TWISST) of the three rooted topologies of *major*, *intermedius* and *bokharensis* (*P. minor* outgroup; rolling mean, *k* = 11 windows). Topo 3 (*intermedius* + *bokharensis*, orange) peaks at the candidate interval, where the species topology (Topo 1) falls to near zero. **(B)** *f_dM_* in windows of 50 SNPs (step 25); dashed line, genome-wide mean (0.086); dotted red line, 99th percentile (0.337); orange points, outlier windows. **(C)** Genome polarization (DIEM) per individual in 50-kb windows (orange, *bokharensis*-like; grey, *major* -like), with the chromosome-wide hybrid index on the left; stars mark Lotfabad individuals Z0016 and Z0072, which carry *bokharensis* mitochondrial haplotypes. The orange band (A, B) and outline (C) mark the 50-kb window (1.475–1.525 Mb) containing *BCO2*. *Alt text:* Three plots along chromosome 24, showing topology weights, *f_dM_* and genome polarization, each with a localized shift towards *bokharensis* ancestry in *intermedius* at the *BCO2* interval.

Topology weighting analyses revealed that the predominant topology along chromosome 24 grouped *intermedius* with *major*, consistent with overall patterns of genomic similarity (Fig. 5A). However, within the differentiated interval, there was a localized increase in support for the topology grouping *intermedius* with *bokharensis*, indicating a shift in genealogical relationships specific to this region.

Across the genome, *f_dM_* values were low but consistently positive (mean = 0.086), indicating low-level genome-wide excess allele sharing between *intermedius* and *bokharensis*, consistent with limited gene flow between them. Against this baseline, *f_dM_*peaked in and just upstream of the candidate interval, where three windows exceeded the genome-wide 99th percentile (*f_dM_* = 0.34–0.50; *z* up to 4.2; Fig. 5B), indicating locally elevated allele sharing between *intermedius* and *bokharensis* relative to *major*.

Genome polarization analyses further supported this result and indicated its direction: ancestry within *intermedius* genomes was predominantly associated with *major* across most of the chromosome but shifted toward *bokharensis* within the same interval (Fig. 5C), consistent with introgression from *bokharensis* into *intermedius*.

### The introgressed region contains the carotenoid-processing gene *BCO2*

To identify genes underlying the differentiated region on chromosome 24, we annotated the candidate interval corresponding to the peak of genetic differentiation and localized introgression. This region spans approximately 34.9 kb (1,490,771–1,525,695 bp) and contains or overlaps seven annotated protein-coding genes: *BCO2, TEX12, LOC107214402, SDHD, NKAPD1, DLAT,* and *DIXDC1* (Fig. 4C; Table S5). Among these, *BCO2* encodes *β*-carotene oxygenase 2, an enzyme involved in carotenoid metabolism.

To assess patterns of coding diversity, we compared synonymous (4-fold degenerate) and nonsynonymous (0-fold degenerate) variation in *BCO2* with that of the pooled neighbouring genes (Table S5) within the differentiated region (Table 1). In *intermedius*, synonymous diversity (*π*_4_ and *θ*_4_) at *BCO2* was comparable to that of the neighbouring genes, indicating no evidence for a local reduction in neutral diversity. In contrast, the ratio of nonsynonymous to synonymous nucleotide diversity (*π*_0_*/π*_4_) was approximately eightfold lower at *BCO2* (0.008) than in the neighbouring genes (0.061), whereas the corresponding difference in Watterson’s estimator (*θ*_0_*/θ*_4_) was comparatively modest (0.079 versus 0.088). In the *major* lineage, synonymous diversity at *BCO2* was lower than in the neighbouring genes for both *π*_4_ and *θ*_4_. Together, these patterns are consistent with stronger evolutionary constraint on the coding sequence of *BCO2* in *intermedius* relative to neighbouring genes.

**Table 1:**
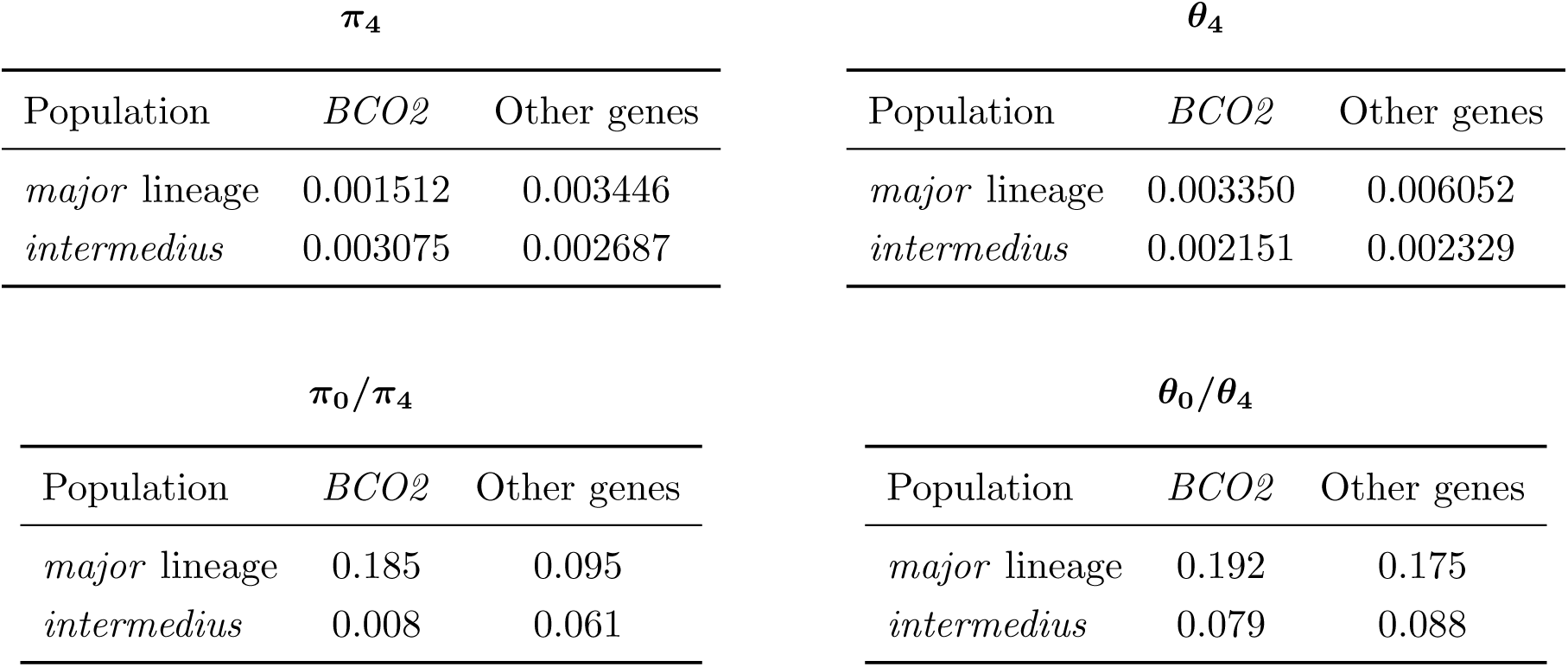
Comparison of synonymous diversity (*π*_4_ and *θ*_4_) and relative coding diversity (*π*_0_*/π*_4_ and *θ*_0_*/θ*_4_) between *BCO2* and the pooled neighbouring candidate genes (*TEX12*, *LOC107214402*, *SDHD*, *NKAPD1*, *DLAT*, and *DIXDC1* ) within the highly differentiated chromosome 24 region. Estimates are based on 42 *major* -lineage (European *major*, *karelini* and *blanfordi* ) and 31 *P. m. intermedius* individuals.

| **π<sub>4</sub>** |  |  | **θ<sub>4</sub>** |  |  |
|---|---|---|---|---|---|
| Population | *BCO2* | Other genes | Population | *BCO2* | Other genes |
| *major* lineage | 0.001512 | 0.003446 | *major* lineage | 0.003350 | 0.006052 |
| *intermedius* | 0.003075 | 0.002687 | *intermedius* | 0.002151 | 0.002329 |
| **π<sub>0</sub>/π<sub>4</sub>** |  |  | **θ<sub>0</sub>/θ<sub>4</sub>** |  |  |
| Population | *BCO2* | Other genes | Population | *BCO2* | Other genes |
| *major* lineage | 0.185 | 0.095 | *major* lineage | 0.192 | 0.175 |
| *intermedius* | 0.008 | 0.061 | *intermedius* | 0.079 | 0.088 |

In contrast to the limited coding differentiation, two highly differentiated SNPs (*F*_ST_ *>* 0.9) were located within the 5*^′^* untranslated region of *BCO2* (NC 031792.1:1495013 G/A and NC 031792.1:1494978 A/G), suggesting that regulatory variation may contribute to the observed phenotypic differences. Across *BCO2*, *intermedius* carried a *bokharensis*-like haplotype, with highly differentiated SNPs and UTR/upstream variants concentrated toward the 5*^′^* end of the gene (Fig. 6). Comparison across representative parid species showed that the full two-SNP combination (A at 1495013, G at 1494978) was observed only in *P. bokharensis* and *P. m. intermedius*, whereas the other examined taxa carried either the *major* combination or other combinations of the two alleles (Table S6). The G allele at 1494978 was also present in the outgroups *Poecile atricapillus* and *Cyanistes caeruleus*, suggesting it represents the ancestral state, whereas the A allele at 1495013 was also found in *P. cinereus*.

**Figure 6:**
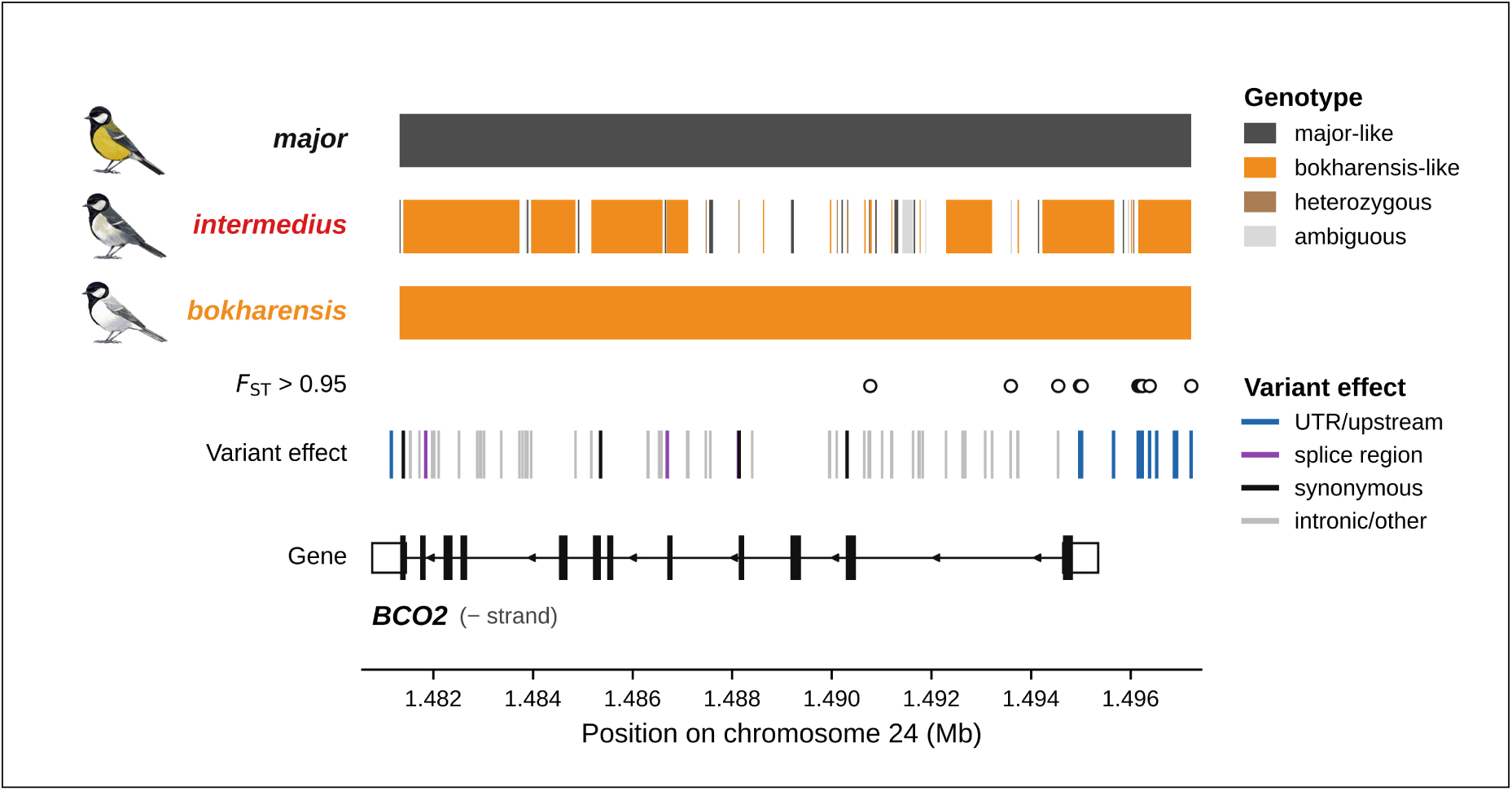
Local haplotype structure and functional annotation across the *BCO2* locus. Informative SNPs across *BCO2* in *intermedius* (Z0046), classified relative to the *major* -lineage (*karelini*, Z0027) and *bokharensis* (B3959) reference genotypes. Orange, *bokharensis*-like; dark grey, *major* -like; brown, heterozygous; light grey, ambiguous. Open circles mark SNPs with *F*_ST_ *>* 0.95, a stricter threshold than that used to define the candidate interval (*F*_ST_ *>* 0.9; Fig. 4C). The variant track shows predicted effects (blue, UTR/upstream; purple, splice region; black, synonymous; grey, intronic or other). The gene model shows the annotated *BCO2* transcript (XM 015649779.3; filled boxes, coding sequence; open boxes, UTRs); arrows indicate transcription on the minus strand. *Alt text:* A SNP-by-SNP haplotype panel across the *BCO2* gene showing *bokharensis*-like alleles in *intermedius*, concentrated towards the 5*^′^* end, with variant annotations and the gene model below.

## Discussion

Whether introgressive hybridization can supply genetic variation to populations at species’ range margins, where adaptation may be constrained by genetic drift and maladaptive gene flow, remains an important evolutionary question. We investigated this question in *P. m. intermedius*, a peripheral great tit population near the contact zone between the Eurasian *major* and Central Asian *bokharensis* lineages. Although genetically differentiated, *intermedius* retains predominantly *major* -lineage ancestry, with limited genomewide introgression from *bokharensis*. Notably, its nucleotide diversity is only slightly lower than that of *major*, contrasting with the genetic erosion expected at range margins. Against this genomic background, one region stands out: a 35-kb segment on chromosome 24 containing the carotenoid-cleaving gene *BCO2*. In this highly differentiated region, allele sharing with *bokharensis* (*f_dM_* ) far exceeds the positive genome-wide mean, suggesting that, although *intermedius* has received *bokharensis* alleles across the genome, they have been preferentially retained in this region, a pattern supported by local genealogical and ancestry analyses. Reduced nucleotide diversity and negative Tajima’s *D* in this region suggest a role for selection, and introgressed variation near *BCO2* may contribute to the distinctive grey plumage of *intermedius*. Together, these findings suggest that introgression may maintain genetic variation in peripheral populations while also supplying locally favoured alleles that can facilitate adaptation at extreme range margins.

### Evolutionary Divergence at the Species’ Range Margin

Peripheral populations are often expected to exhibit elevated genetic differentiation and reduced diversity because of smaller effective population sizes, geographic isolation, and genetic drift (Eckert et al. 2008; Pironon et al. 2017). Consistent with the expectation of increased differentiation, genome-wide *F*_ST_ between *P. m. intermedius* and European *major* reached 0.056, nearly fourfold higher than the mean among European populations (*F*_ST_ = 0.015) and approaching the highest values in Europe, which involve the relatively isolated Mediterranean island populations of Corsica, Sardinia and Crete (Spurgin et al. 2024). Comparable genetic divergence between eastern and western populations in Iran has been reported for several other passerines, coinciding with the region’s zoogeographic boundaries (Javaheri Tehrani et al. 2025). This distinctiveness was further supported by PCA and ADMIXTURE (*K* = 3), in which *intermedius* formed a distinct cluster (Fig. 3A, C). Despite this pronounced differentiation, nucleotide diversity in *intermedius* was only slightly lower than in *major* (Fig. S6B), indicating that differentiation has not been accompanied by substantial genetic erosion. This contrasts with the expectation that peripheral isolation and genetic drift should reduce within-population diversity.

The contrasting allele-frequency spectra of *intermedius* and European *major* provide further insight into their distinct evolutionary histories. In European *major*, predominantly negative Tajima’s *D* indicates an excess of rare variants, consistent with demographic expansion. Our MSMC2 reconstruction independently reveals historical growth in effective population size (Fig. S3), placing this pattern within a broader history of population expansion documented by previous PSMC analyses (Laine et al. 2016), mitochondrial and genomic evidence of postglacial recolonization of Europe from a southeastern refugium (Kvist et al. 1999; Spurgin et al. 2024), and demographic reconstructions across the great tit complex (Song et al. 2020). In contrast, predominantly positive Tajima’s *D* in *intermedius* (Fig. S6A) indicates a relative deficit of rare variants, suggesting a different demographic or population history from European *major*. Although population contraction and drift could contribute to this pattern, drift alone would also be expected to reduce nucleotide diversity, whereas population structure or admixture can shift the allele-frequency spectrum while maintaining diversity.

### Genome-Wide Introgression and a Localized Ancestry Shift

The intermediate geographic position and phenotypic resemblance of *intermedius* to *bokharensis* have long suggested possible gene flow between the *major* and *bokharensis* lineages (Harrap and Quinn 1996; Martens 1996), but this hypothesis has not previously been tested with whole-genome data. Our genome-wide analyses now provide this evidence, revealing that gene flow from the *bokharensis* lineage has contributed to the ancestry of *intermedius*. Most *intermedius* individuals carry only limited *bokharensis* ancestry, with hybrid indices and interclass heterozygosity consistent with later-generation admixture. Only three individuals exhibited genomic patterns characteristic of putative early-generation hybrids (Fig. 3B). Together with increasing *bokharensis*-associated ancestry towards the range of *bokharensis* (Fig. 3C) and mitochondrial haplotype sharing across lineage boundaries (Fig. 2A; Fig. S2), these findings support gene flow where the two lineages meet. This gene flow may also explain the discordance between nuclear and mitochondrial phylogenies. Our mitogenome tree was consistent with previous mitochondrial phylogenies (Kvist et al. 2003; Song et al. 2020), recovering *bokharensis* as sister to *major*, whereas the nuclear species tree placed *bokharensis* as sister to *cinereus* (Fig. 2B). Such mitonuclear discordance is common in animals and particularly in birds, where it frequently reflects introgression (Toews and Brelsford 2012; Toews et al. 2014; Alaei Kakhki et al. 2023). The internal branch uniting *bokharensis* and *cinereus* also showed the highest gene-tree discordance in the species tree, with only 43% of quartets supporting it and an excess of one alternative topology (q2 = 0.36 vs q3 = 0.21). Incomplete lineage sorting during the rapid Pleistocene diversification of the complex may contribute to this discordance, but would produce both alternative topologies at similar frequencies; genome-wide tests of excess allele sharing indicate that introgression also plays a role: *f* -branch statistics revealed excess allele sharing between *bokharensis* and all lineages of the *major* clade, with the strongest signal involving *intermedius* (*f_b_ ≈* 0.20; Fig. 2C). Mitochondrial introgression between *bokharensis* and *major* would thus account for their sister relationship in the mitogenome tree, while nuclear gene flow would explain the elevated gene-tree discordance at this internal branch. The predominance of *major* -lineage ancestry in *intermedius* may reflect the progressive dilution of introgressed ancestry through continued connectivity with core *major* -lineage populations, and potentially selection against deleterious introgressed variation (Harris and Nielsen 2016; Juric et al. 2016; Veller et al. 2023). The observed introgression is consistent with expectations for range expansions, under which even limited interbreeding leads to asymmetric introgression from the resident into the expanding population (Currat et al. 2008). MSMC2 reconstructions showing long-term growth of the *major* lineage, in contrast to a small and stable *bokharensis* (Fig. S3), identify *major* as the expanding and *bokharensis* as the resident lineage, so introgression is expected predominantly from *bokharensis* into *major* -lineage populations such as *intermedius*. Because such introgression can occur at neutral loci without selection, introgressed ancestry does not directly imply adaptation, although it can increase nucleotide diversity and thereby the adaptive potential of the recipient population. We therefore used the genome-wide *f*_dM_ distribution, with its positive mean, as a baseline, treating regions far above it as candidates for adaptive introgression.

Against this predominantly *major* -lineage genomic background, a localized region on chromosome 24 exhibits pronounced *bokharensis*-associated ancestry. Window-based *F*_ST_ between *intermedius* and *major* reached approximately 0.65 in this region, more than tenfold the genome-wide average (Fig. 4). Excess allele sharing between *intermedius* and *bokharensis* (*f*_dM_), increased support for local genealogies grouping these lineages, and genome polarization all point to gene flow from *bokharensis* into *intermedius* in this region (Fig. 5). Such a heterogeneous genomic landscape of introgression is widely observed and reflects interactions among selection, recombination, and genomic architecture (Gaczorek et al. 2024; Justen et al. 2025). Notably, chromosome 24 is a microchromosome. Avian microchromosomes generally exhibit elevated recombination rates (Ellegren et al. 2012; Gossmann et al. 2014; Stapley et al. 2017; Blain et al. 2025), which may facilitate introgression by uncoupling donor alleles from linked deleterious or incompatible variants (Juric et al. 2016; Schumer et al. 2018). The location of the candidate region on chromosome 24 therefore raises the possibility that its genomic context has facilitated the persistence of *bokharensis*-associated ancestry, although local recombination rates have not been estimated. Moreover, several features of this *bokharensis*-introgressed region on chromosome 24 are consistent with positive selection: reduced nucleotide diversity, most pronounced in *intermedius*, a shift towards negative Tajima’s *D* (Fig. 4B), and the retention of *bokharensis* ancestry here, unlike most of the genome.

### Introgression as a Potential Source of Adaptive Variation at Range Margins

The introgressed interval on chromosome 24 contains or overlaps seven genes (Table S5), of which *BCO2*, encoding a carotenoid-cleaving enzyme, is the only one with an established role in carotenoid metabolism and pigmentation (Walsh et al. 2012). Two highly differentiated variants in its 5*^′^* untranslated region form a two-SNP combination shared by *intermedius* and *bokharensis* but absent from the other sampled parids (Table S6). No amino acid substitution was fixed between *intermedius* and *major*, and the ratio of nonsynonymous to synonymous diversity at *BCO2* was approximately eightfold lower in *intermedius* than in neighbouring genes (Table 1). Together, these patterns raise the hypothesis that regulatory rather than protein-coding variation at *BCO2* contributes to the grey plumage of *intermedius*, although the functional effects of the two 5*^′^*UTR variants remain untested. Regulatory changes are common in pigmentation evolution (Stern and Orgogozo 2008), including at *BCO2* in *Vermivora* warblers (Toews et al. 2016) and *Manacus* manakins (Lim et al. 2024) and at *ASIP* in wheatears (Lutgen et al. 2025), and in domestic chickens the yellow-skin phenotype is caused by regulatory *BCO2* variation introgressed from the grey junglefowl (Eriksson et al. 2008). Because *BCO2* degrades carotenoids, one prediction is increased expression in developing feathers of individuals carrying the introgressed haplotype, reducing carotenoid deposition; alternatively, the variants may alter the timing or tissue specificity of expression. Expression analyses and functional assays are needed to test these possibilities.

Why such variation might be favoured in *intermedius* remains uncertain, but the similar arid habitats occupied by *intermedius* and *bokharensis* could explain it: alleles shaped by selection in *bokharensis* may be advantageous in *intermedius* under comparable ecological conditions. Yellow breast coloration in great tits largely reflects dietary carotenoids obtained from caterpillars (Slagsvold and Lifjeld 1985; Eeva et al. 2010), although it also has a genetic component (Fitze et al. 2003). To our knowledge, *BCO2* is one of the first candidate loci proposed for population-level differences in carotenoid-based plumage in this species. *P. m. intermedius* inhabits the arid northeast of Iran, where the lepidopteran fauna is comparatively species-poor (Noori et al. 2024; Noori et al. 2025). If carotenoid-rich prey are less available in these habitats, variation affecting carotenoid deposition could alter the dependence of plumage coloration on dietary resources. Alternatively, greyer plumage might improve background matching in open, sparsely vegetated environments, whereas yellow plumage has been proposed to match tree foliage (Slagsvold and Lifjeld 1985). Although dietary carotenoid availability, feather pigmentation, and the camouflage value of plumage coloration were not directly assessed, both hypotheses would explain why *bokharensis*-derived variation at *BCO2* has been retained in *intermedius*.

Together, our findings suggest that introgression may contribute to genomic and potentially phenotypic divergence at a species’ range margin without substantially altering the recipient population’s overall ancestry. Although genome-wide admixture between *intermedius* and *bokharensis* is limited, a localized region on chromosome 24 exhibits a pronounced introgression signal and contains candidate regulatory variation at *BCO2*. Introgression may therefore help peripheral populations in two complementary ways: low-level genetic exchange across the genome can introduce variation and buffer against its loss through drift, whereas particular introgressed variants may facilitate local adaptation if they confer an advantage in peripheral environments. These findings connect to broader hypotheses about the role of introgression in population persistence and evolutionary rescue (Bell and Gonzalez 2009; Carlson et al. 2014), exemplified by introgression-driven adaptation to extreme pollution in Gulf killifish (Oziolor et al. 2019), particularly as climate change shifts species ranges and creates new opportunities for contact between divergent lineages (Chunco 2014; Brauer et al. 2023). Our study thus identifies a potential route through which hybridization may contribute to evolutionary diversification, local adaptation, and the persistence of peripheral populations.

## Materials and Methods

### Genomic sampling

To investigate the evolutionary origin of *P. m. intermedius* and its relationship to neighbouring *major* and *bokharensis* populations, we generated whole-genome resequencing data for 63 Iranian individuals representing three subspecies: *P. m. intermedius* (*n* = 31), *P. m. karelini* (*n* = 23), and *P. m. blanfordi* (*n* = 9) (Dickinson and Christidis 2014). Sampling of *P. m. intermedius* spanned northeastern Iran and included sites near the Turkmenistan border (Lotfabad; Fig. 1A), corresponding to the putative contact zone between the *major* and *bokharensis* lineages. To place these populations within a broader phylogenomic context, we additionally included seven *bokharensis* individuals from Kazakhstan and Kyrgyzstan obtained from the Senckenberg Natural History Collections Dresden (SNSD), three *cinereus* individuals from Nepal obtained from SNSD, 10 publicly available *major* genomes and 29 publicly available *minor* genomes downloaded from the NCBI Sequence Read Archive (SRA). Sampling locations are shown in Fig. 1A, with the Iranian sampling localities shown in the right-hand map and inset. A complete list of samples, accession numbers, and metadata is provided in Table S1. Because analyses differed in their data requirements, each was performed on a subset of individuals selected according to analysis-specific criteria (Table S2). Throughout the manuscript, we use the term “lineage” to refer to genetically defined population clusters within the *P. major* species complex and closely related taxa without implying formal taxonomic rank.

Genomic DNA was extracted from blood samples stored in Queen’s Lysis Buffer or tissue samples preserved in 96% ethanol using the DNeasy Blood and Tissue Kit (Qiagen) following the manufacturer’s protocol. DNA from frozen museum tissues was extracted using either the Promega Wizard Genomic DNA Purification Kit or the Qiagen MagAttract HMW DNA Kit according to the manufacturers’ instructions. DNA concentration and quality were assessed using a Qubit Fluorometer (dsDNA Broad Range Assay; Thermo Fisher Scientific) and a NanoDrop spectrophotometer. Libraries were sequenced as 150-bp paired-end reads on Illumina NovaSeq 6000 platforms, targeting a mean coverage of approximately 32X (range: 16–142X; Table S1).

### Mapping and variant calling

Raw sequencing reads were assessed for quality using FastQC v0.12.1. Adapter sequences and low-quality bases were removed using Trimmomatic v0.39 (Bolger et al. 2014). Filtered reads were aligned to the *Parus major* reference genome (GenBank accession GCF 001522545.3) using BWA-MEM2 v2.2.1 with default parameters. Resulting SAM files were converted to sorted and indexed BAM files using SAMtools v1.2 (Li et al. 2009). Duplicate reads were identified and marked using MarkDuplicatesSpark implemented in GATK v4.5.0 (Van der Auwera and O’Connor 2020).

Base Quality Score Recalibration (BQSR) was performed following GATK Best Practices pipeline (McKenna et al. 2010), using a high-confidence set of SNPs as known sites. Variants were called for each individual using HaplotypeCaller in GVCF mode on base-quality recalibrated BAM files. Individual GVCF files were combined using CombineGVCFs, followed by joint genotyping of variant and invariant sites using GenotypeGVCFs with the --include-non-variant-sites flag.

### Data filtering

To minimize biases associated with repetitive genomic regions, we excluded all sites overlapping annotated repeat regions in the *Parus major* reference genome from the multi-sample VCF file using BCFtools (Danecek et al. 2021). Variant calls generated separately for each chromosome were subsequently merged into a single genome-wide VCF file. We then applied a series of filters to retain high-quality biallelic SNPs. Sites overlapping indels or located within 10 bp of indels were removed, and variants with mapping quality *<* 30 were excluded. Additional filtering was performed using VCFtools (Danecek et al. 2011) to retain SNPs with a minimum read depth of 5, minor allele frequency *≥* 0.05, minimum site quality *≥* 30, minimum genotype quality *≥* 20, and a maximum missingness of 20% per site. Unless otherwise specified, all downstream population genomic analyses were conducted using this final filtered SNP dataset.

### Sex determination and relatedness filtering

To assess and account for potential effects of sex-specific genomic differentiation (sex bias) (Chen et al. 2025), and to avoid non-independence among samples, we inferred genetic sex and filtered closely related individuals prior to subsequent analyses. Genetic sex was inferred using a coverage-based approach that exploits differences in read depth between sex chromosomes and autosomes. Sequencing depth was calculated using SAMtools depth v1.13 across three genomic regions: the Z chromosome (NC 031799.1), an autosomal reference chromosome (chromosome 3; NC 031770.1), and the mitochondrial genome (NC 040875.1). For each individual, mean coverage was calculated for each region, and the ratio of Z-chromosome to autosomal coverage was used to infer sex. In birds with a ZW sex-determination system, this ratio is expected to be close to 1 in genetic males (ZZ) and approximately 0.5 in genetic females (ZW). Individuals with a Z-to-autosome coverage ratio *<* 0.75 were classified as females, whereas those with ratios *≥* 0.75 were classified as males. Mitochondrial coverage was used as an internal control to verify sequencing consistency. Sex assignments for all individuals are summarized in Table S1. Pairwise relatedness among individuals was estimated using NGSrelate v2 (Korneliussen and Moltke 2015). To minimize the influence of close kinship on downstream analyses, one individual from each pair with a relatedness coefficient *>* 0.25 was removed. All subsequent nuclear analyses were conducted using the resulting set of unrelated individuals (*n* = 112); individuals removed by this filter were retained for mitogenome analyses (Table S1).

### Mitochondrial genome phylogeny and haplotype analysis

To resolve the phylogenetic relationships within the *P. major* complex and determine the evolutionary placement of *intermedius* populations, we reconstructed mitochondrial phylogenies and haplotype networks. Complete mitochondrial genomes were extracted and *de novo* assembled from whole-genome sequencing (WGS) data for 99 individuals using GetOrganelle (Jin et al. 2020), utilizing the published *Parus major* mitogenome (GenBank: NC 026293.1) as a reference for assembly verification. The resulting mitogenomes were aligned via the MAFFT algorithm within the UGENE v42.0 environment (Okonechnikov et al. 2012). Maximum-likelihood (ML) phylogenetic inference was performed using IQ-TREE v2.1.2 (Minh et al. 2020) under the TIM2+F+I substitution model, selected as best-fitting by ModelFinder according to the Bayesian information criterion (Kalyaanamoorthy et al. 2017). Node support was evaluated using 1,000 ultrafast bootstrap replicates, and the blue tit (*Cyanistes caeruleus*) was designated as the outgroup to root the tree. To visualize fine-scale maternal relationships, a mitochondrial haplotype network was constructed in PopART (Leigh and Bryant 2015) using the TCS network method with a 95% connection limit to identify genealogical connections across the complex.

### Nuclear genome phylogeny and species tree inference

To reconstruct phylogenetic relationships across the nuclear genome, assess the mitogenome discordance, and determine the placement of *intermedius* relative to other taxa within the *P. major* complex, we reconstructed species trees from whole-genome data. To minimize potential biases associated with sex-linked genomic variation and uneven sampling among populations, analyses were restricted to one or two genetically representative male individuals per population. For phylogenomic analyses incorporating both variant and invariant sites, the nuclear genome was partitioned into non-overlapping 10-kb loci, and a maximum-likelihood gene tree was inferred for each locus.

To evaluate the sensitivity of phylogenetic inference to data filtering, we generated four sets of FASTA alignments using different thresholds for minimum read depth (DP), minimum window coverage (PW), and maximum missing data across individuals (MD). The filtering strategies were: (1) DP = 1, PW = 50%, MD = 15%; (2) DP = 5, PW = 50%, MD = 15%; (3) DP = 1, PW = 50%, MD = 5%; and (4) DP = 1, PW = 80%, MD = 10% (Table S3). Gene trees were inferred for each filtered dataset using IQ-TREE v2.1.2 under the GTR+I+G substitution model with 1,000 ultrafast bootstrap replicates. As all filtering strategies recovered concordant species-level topologies, subsequent analyses were conducted using the dataset filtered with DP = 5, PW = 50%, and MD = 15%.

To minimize the influence of intralocus recombination on species tree inference, each locus was tested for evidence of recombination using the Pairwise Homoplasy Index (Φ*_w_*) implemented in PhiPack (Bruen et al. 2006), using the default sliding window size (100 bp) and 1,000 permutations. Loci showing significant evidence of recombination (Φ*_w_* permutation test, *P ≤* 0.05) were excluded from downstream analyses. To further reduce the effects of physical linkage, the remaining loci were pruned by retaining only windows separated by at least 10 kb on the same chromosome. This procedure reduced the dataset from 90,549 initial loci to 35,097 recombination-free and linkage-pruned gene trees.

Species trees were inferred under the multispecies coalescent using ASTRAL v5.7.8 (Zhang et al. 2018); branch support was assessed using local posterior probabilities (LPP), and quartet support (q1–q3) was used to quantify gene-tree discordance for each internal branch.

### Genome-wide tests for introgression

To assess genome-wide patterns of introgression among lineages, we used Dsuite (Malinsky et al. 2021) to estimate Patterson’s *D* statistic (ABBA–BABA test) (Durand et al. 2011) and the *f*_4_-ratio for all population trios, with significance assessed by block-jackknife resampling. Analyses were based on the repeat-masked autosomal SNP dataset described above, excluding indels, sites lacking an alternative allele, sites fixed for the alternative allele and sites with more than 20% missing data, and thinned to one SNP per 100 bp (approximately 5.9 million SNPs). Analyses were conducted on the same 12 high-coverage male individuals used for species tree inference (one or two per lineage; Table S2), selected to avoid individuals with evidence of recent admixture, with the blue tit (*Cyanistes caeruleus*) as outgroup. To place excess allele sharing in a phylogenetic context, *f*_4_-ratio results were summarized as *f* -branch (*f_b_*) statistics (**Malinsky2018**) mapped onto the ASTRAL species tree (Fig. 2C).

### Demographic inference using MSMC2

To infer historical changes in effective population size (*N_e_*) of the *major* and *bokharensis* lineages, we performed coalescent-based analyses using MSMC2 (Multiple Sequentially Markovian Coalescent; Schiffels and Durbin 2014; Schiffels and Wang 2020). Because MSMC2 is sensitive to sequencing depth, analyses were restricted to high-coverage (*>*20*×*) male individuals; males were used because they are the homogametic sex (ZZ) in birds, which avoids biases associated with the hemizygous Z chromosome in females (Chen et al. 2025). For each lineage, a single high-coverage male genome was selected (*n* = 1 per lineage), as MSMC2 can infer long-term demographic trajectories from a single diploid genome. Individuals from the *minor* lineage were excluded because their coverage (4.07–16.45*×*) was below the threshold required for reliable inference. Individual-specific VCF and mask files were generated from recalibrated BAM files using bamCaller.py from the msmc-tools package, and MSMC2 was run separately for each lineage with default time-interval settings. Coalescent time and *N_e_* were scaled to real units using a mutation rate of 2.0 *×* 10*^−^*^9^ substitutions per site per generation and a generation time of two years (Laine et al. 2016).

### Population genomic structure and hybrid ancestry

For population structure analyses, including principal component analysis (PCA) and ADMIXTURE, we used autosomal biallelic single-nucleotide polymorphisms (SNPs) derived from the filtered dataset described above. Biallelic SNPs were retained using BCFtools (Danecek et al. 2021). To ensure the use of approximately independent SNPs, variants were thinned by retaining one SNP per 1 kb across the genome, thereby reducing the effects of linkage disequilibrium.

PCA was performed with the SNPRelate package v1.30.0 in R (Zheng et al. 2012) after converting the data to Genomic Data Structure (GDS) format, using the snpgdsPCA function. A range-wide PCA was used to place *intermedius* among all lineages of the complex. For this analysis, individuals with more than 10% missing genotypes were excluded (Table S2). Because the deep divergence between *minor* and all other lineages accounted for most of the variance captured by PC1 and compressed the structure among the remaining lineages, the analysis was repeated after excluding *minor* (Fig. S4). A regional PCA of *major*, *intermedius* and *bokharensis* (48 individuals) was performed to examine structure among the three focal lineages (Fig. 3A).

For ADMIXTURE analyses, genotype data were converted from VCF to PLINK binary format. Individual ancestry proportions were inferred using the maximum-likelihood clustering algorithm implemented in ADMIXTURE v1.3.0 (Alexander et al. 2009). Analyses were conducted assuming *K* = 2–4 ancestral clusters, with 10 independent replicate runs performed for each value of *K* to assess convergence across runs. Cross-validation error was lowest at *K* = 2 (Table S4), which captured the primary split between the *major* and *bokharensis* lineages; we also present *K* = 3, at which an additional component specific to *intermedius* emerged, and evaluated support across values of *K* using the Evanno Δ*K* method (Evanno et al. 2005) implemented in CLUMPAK (Kopelman et al. 2015). Individuals were assigned to a genetic cluster if their inferred ancestry proportion (*Q*-value) exceeded 0.9, whereas individuals with lower values were classified as admixed.

To further quantify hybrid ancestry and assign hybrid classes of *intermedius* lineage, we estimated hybrid index values and interspecific heterozygosity using the triangulaR package (Wiens et al. 2025) implemented in R. The hybrid index was defined as the proportion of alleles inherited from the two parental lineages: including *major* and *bokharensis*. We used the VCF file including only variant sites. Hybrid index and interspecific heterozygosity estimates were used to visualize the distribution of individuals in hybrid index–heterozygosity space and to classify individuals into parental, early-generation hybrid, and backcross categories following established expectations.

### Genome-wide differentiation and diversity analyses

To identify genomic regions associated with differentiation and characterize patterns of genetic diversity, we adopted a stepwise analytical framework combining genome-wide summaries and targeted analyses of candidate regions.

First, genome-wide differentiation between *intermedius* and European *major* was quantified using Weir and Cockerham’s *F*_ST_ as implemented in VCFtools v0.1.17, using the repeat-masked autosomal SNP dataset described above.

We then focused on three focal lineages (*major*, *intermedius*, and *bokharensis*) to characterize genome-wide patterns of differentiation and diversity. Sliding-window scans of *F*_ST_ were performed between *major* and *intermedius*, as well as between *intermedius* and *bokharensis*, using 50 kb windows with a 25 kb step size. Genome-wide nucleotide diversity (*π*) and Tajima’s *D* were calculated separately for each lineage using scikit-allel v1.3.13 (Miles et al. 2020), based on biallelic SNPs without applying a minor allele frequency filter to avoid biasing site-frequency spectrum–based estimates. Windows with low numbers of segregating sites were excluded to reduce stochastic variance.

Genome-wide scans identified multiple candidate regions showing elevated differentiation between *major* and *intermedius*. Regions were prioritized for further analysis based on concordant patterns across complementary statistics, including *F*_ST_, nucleotide diversity (*π*), Tajima’s *D*, and absolute sequence divergence (*d_XY_* ). Chromosome 24 showed the strongest concordant signal across these statistics and was therefore selected for detailed downstream analyses.

Fine-scale sliding-window analyses across chromosome 24 were then performed to further characterize local patterns of differentiation and diversity. Absolute sequence divergence (*d_XY_* ) between populations was estimated using Pixy (Korunes and Samuk 2021), which incorporates both variant and invariant sites to provide unbiased estimates. To further examine differentiation at the candidate locus, we conducted a SNP-level analysis across the genomic interval spanning approximately 1.28–1.70 Mb on chromosome 24. Per-site *F*_ST_ values between *major* and *intermedius* were calculated using the Weir and Cockerham estimator implemented in VCFtools. SNP-level *F*_ST_ values were visualized along genomic coordinates using ggplot2 in R to identify localized peaks of differentiation. Highly differentiated SNPs were defined as sites with *F*_ST_ *>* 0.9.

To provide gene-level context for highly differentiated sites, gene annotations for the candidate region were obtained from the *P. major* genome annotation, and overlaps between highly differentiated SNPs and annotated genes were identified using BEDTools.

### Localized introgression analyses

For localized introgression analyses, we focused on the same set of individuals representing the three focal lineages (*major*, *intermedius*, and *bokharensis*) used in the genome-wide differentiation analyses. To test whether genomic differentiation on chromosome 24 is associated with introgression among lineages, we applied three complementary approaches to detect, quantify, and interpret introgressed regions: the *f_dM_*statistic, topology weighting using TWISST, and genome polarization using DIEM.

To identify regions of excess allele sharing, we calculated *f_dM_* using Dsuite (Dinvestigate; Malinsky et al. 2021) with P1 = *major* (*n* = 9), P2 = *intermedius* (*n* = 29), P3 = *bokharensis* (*n* = 5), and one high-coverage *P. minor* as outgroup. After masking genotypes with depth *<*5 and retaining biallelic SNPs with *≤*20% missing data, *f_dM_* was estimated across all autosomes in windows of 50 SNPs (step 25 SNPs). We used the genome-wide mean *f_dM_* as a baseline (0.086, SD = 0.098), considered windows above the genome-wide 99th percentile (*f_dM_ >* 0.337) as outliers, and report each window’s deviation from the mean as a *z*-score.

To assess whether local genealogical relationships along chromosome 24 are consistent with introgression, we applied topology weighting using TWISST (Martin and Van Belleghem 2017). The chromosome-specific dataset was phased using Beagle v5.5. Sliding windows of 500 SNPs with a step size of 100 SNPs were defined, and maximum-likelihood trees were inferred for each window using IQ-TREE.

TWISST was used to quantify the relative support for alternative rooted topologies among *major*, *intermedius*, and *bokharensis*, with a single *Parus minor* individual included as an outgroup.

To quantify ancestry composition and direction of gene flow along chromosome 24, we performed genome polarization using DIEM (Baird et al. 2023). VCF files were converted to DIEM input format using the diemr R package. Parental populations were defined *a priori* based on population structure analyses, excluding admixed individuals from parental allele frequency estimation.

### Annotation of candidate loci for adaptation

To identify candidate genes potentially underlying adaptive divergence between *P. m. major* and *P. m. intermedius*, we focused on chromosome 24, which showed a strong and consistent signal of elevated differentiation across multiple analyses (*F*_ST_, *π*, Tajima’s *D*, and *d_XY_* ). Candidate regions were first located by identifying windows within the top 1% of the empirical genome-wide *F*_ST_ distribution; outlier windows on chromosome 24 were merged into a contiguous region of elevated differentiation. Within this region, the candidate interval was then defined by the cluster of highly differentiated SNPs (*F*_ST_ *>* 0.9; see above), spanning 1,490,771–1,525,695 bp. To identify candidate genes, this interval was intersected with the *Parus major* reference genome annotation (GFF3) using BEDTools, and genes overlapping the interval were considered candidate loci.

### Synonymous and nonsynonymous diversity

To compare coding diversity among genes within the differentiated region, we identified four-fold and zerofold degenerate coding sites for each annotated gene (*BCO2*, *TEX12*, *LOC107214402*, *SDHD*, *NKAPD1*, *DLAT*, and *DIXDC1* ) using the *Parus major* reference genome annotation. Degeneracy classes were assigned from the longest annotated coding transcript for each gene, and genomic coordinates of four-fold and zero-fold degenerate sites were extracted.

For each population (*major* lineage, *n* = 42: European *major*, *karelini* and *blanfordi* ; *P. m. intermedius*, *n* = 31), genotype likelihoods were estimated in ANGSD using the reference genome as both the reference and ancestral sequence. Site frequency spectra were inferred separately for four-fold and zero-fold degenerate sites using realSFS. Nucleotide diversity (*π*) and Watterson’s estimator (*θ_W_* ) were calculated directly from the inferred site frequency spectra, and values were normalized by the number of informative sites. To assess the relative strength of constraint on coding sequence, we calculated the ratios *π*_0_*/π*_4_ and *θ*_0_*/θ*_4_ for each gene. For comparison with *BCO2*, estimates from the remaining six genes were pooled to obtain regional background values.

### Comparative analysis of the *BCO2* 5*^′^* UTR haplotype

To assess the evolutionary distribution of the differentiated *BCO2* 5*^′^* untranslated region haplotype, homologous genomic sequences spanning the two highly differentiated SNPs were extracted from publicly available genome assemblies of representative parid species. Orthologous regions were identified using whole-genome alignments and manually inspected to confirm synteny. Allelic states at positions homologous to NC 031792.1:1495013 and NC 031792.1:1494978 were compared across species, and haplotypes were classified according to the combination of alleles present. Species lacking a complete sequence across the region were scored as partial or missing. Haplotype classifications for all examined species are summarized in Table S6.

## Author contributions

S.J.T. led sample collection, laboratory work, computational analysis and interpretation, and manuscript writing. N.A.K. developed analytical scripts, contributed to the interpretation of results, and provided strategic and supervisory input throughout the analytical phases of the study. Y.-C.C. performed laboratory work, generated recalibrated BAM files, and conducted preliminary computational analyses.

M.P. provided samples from collections at SNSD (whole skins and frozen blood [coll. J. Martens]). M.A. contributed substantially to project design and planning and coordinated and carried out sample collection in Iran. K.v.O. contributed to the conceptual development, project design, and early-stage planning of the study. M.B. contributed to the early conceptual framework, particularly regarding hybrid zone dynamics, and provided input during the initial planning phase. T.I.G. conceived the study, supervised the project, contributed to analysis and secured funding. All authors contributed to manuscript revision and approved the final version.

## Acknowledgements

We gratefully acknowledge the Senckenberg Natural History Collections Dresden for providing samples used in this research. We thank Tooba Mohammadian, Yahya Tarahomi, and Ayda Rastegar for assistance with fieldwork, and Ferdowsi University of Mashhad for logistical and technical support. We are grateful to members of CSB lab for helpful discussions and constructive comments on earlier versions of the manuscript and technical support.

## Data accessibility

All newly generated sequencing data will be deposited in the National Center for Biotechnology Information (NCBI) Sequence Read Archive (SRA) upon acceptance of the manuscript, and the filtered VCF files used in all analyses will be made available in a public repository (e.g. Dryad). All analysis scripts and computational pipelines used in this study are publicly available on GitHub at https://github.com/tgossmann/Pmajor_Hybrids.

