## Supplemental Information for "Introgressive hybridization as a potential source of evolutionary rescue in an extreme peripheral environment"

### Supplementary Figures

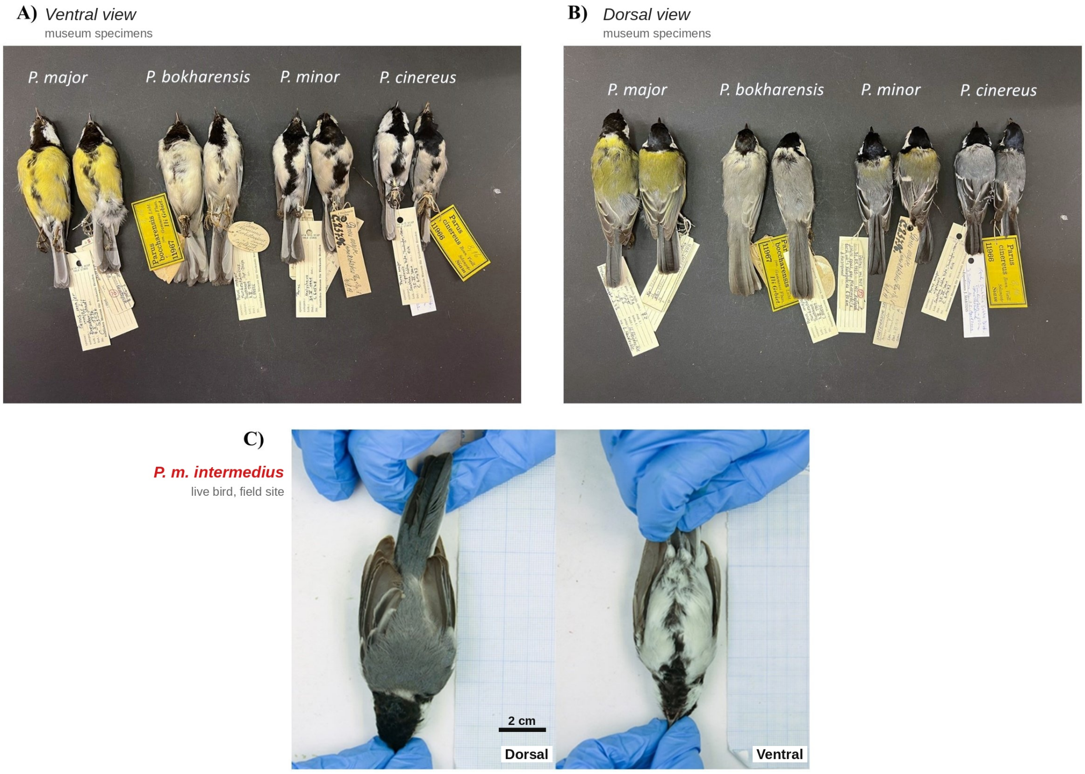


**Figure S1: Variation in plumage coloration across the *P. major* complex.** **(A)** Ventral and **(B)** dorsal views of museum specimens of *P. major*, *P. bokharensis*, *P. minor* and *P. cinereus* (Senckenberg Natural History Collections Dresden). **(C)** Dorsal and ventral views of a live *P. m. intermedius* individual sampled in Lotfabad; the scale bar (2 cm) is approximate, derived from the background graph paper. Photographs were not colour-standardized.

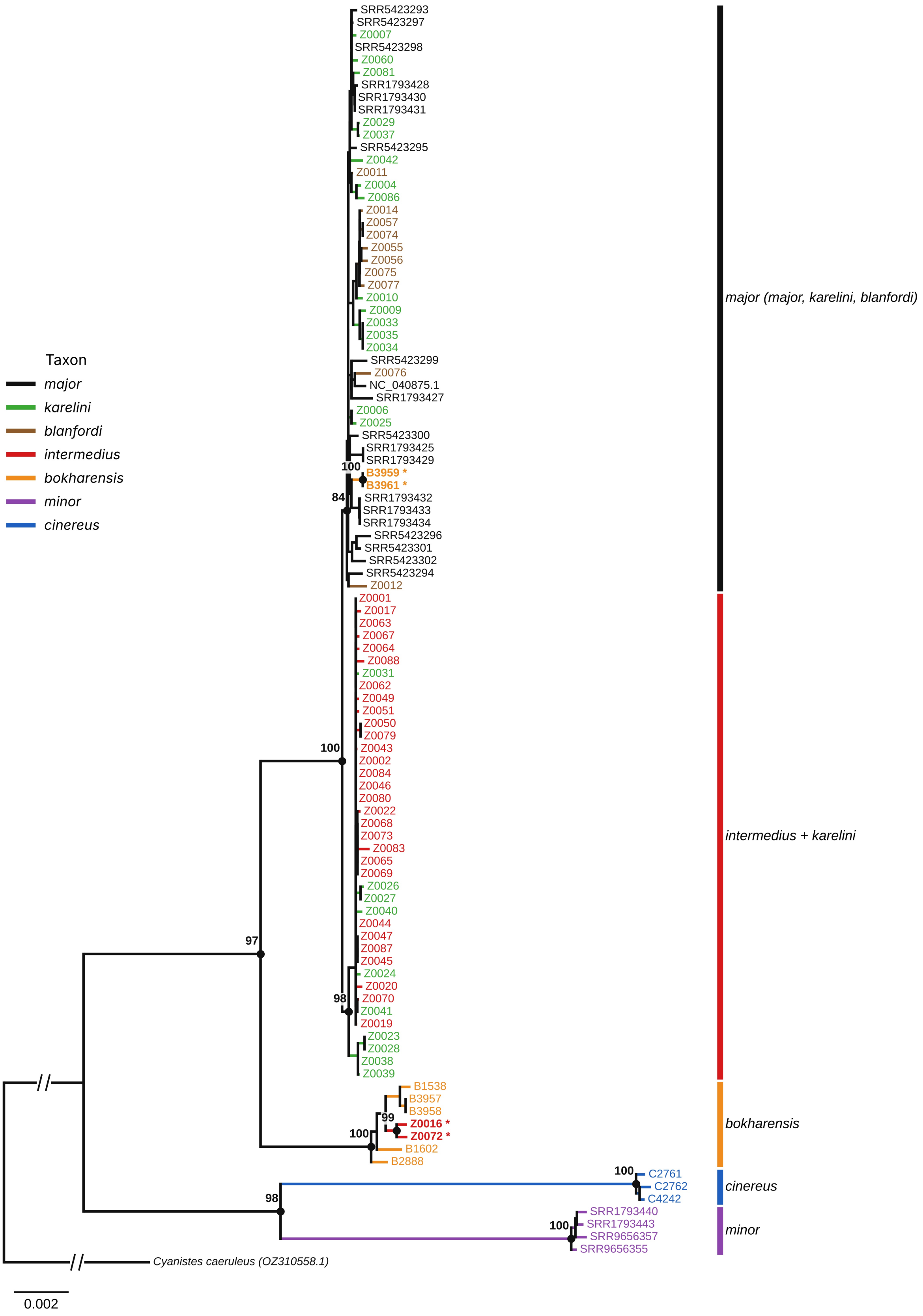


**Figure S2: Maximum-likelihood phylogeny of whole mitochondrial genomes.** IQ-TREE tree (TIM2+F+I) rooted with *Cyanistes caeruleus*; node values are ultrafast bootstrap support (1,000 replicates). Asterisks mark individuals with discordant mitochondrial haplotypes (Z0016, Z0072 within *bokharensis*; B3959, B3961 within *major*). The outgroup branch is shortened (//); scale bar, substitutions per site.

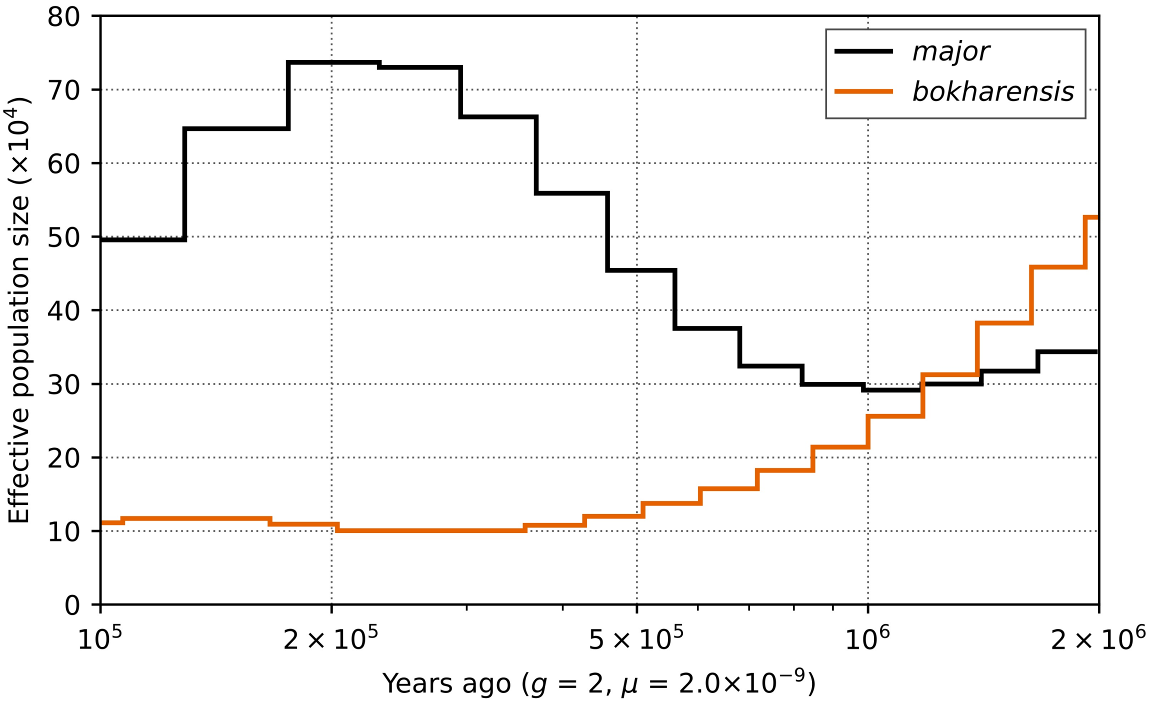


**Figure S3: Historical effective population size inferred using MSMC2.** Changes in effective population size ($N_e$) through time for the *major* (black) and *bokharensis* (orange) lineages, each inferred from a single high-coverage male genome. The *minor* lineage was excluded owing to insufficient sequencing coverage. Time and $N_e$ were scaled using a mutation rate of $2.0 \times 10^{-9}$ substitutions per site per generation and a generation time of two years (Laine et al. 2016).

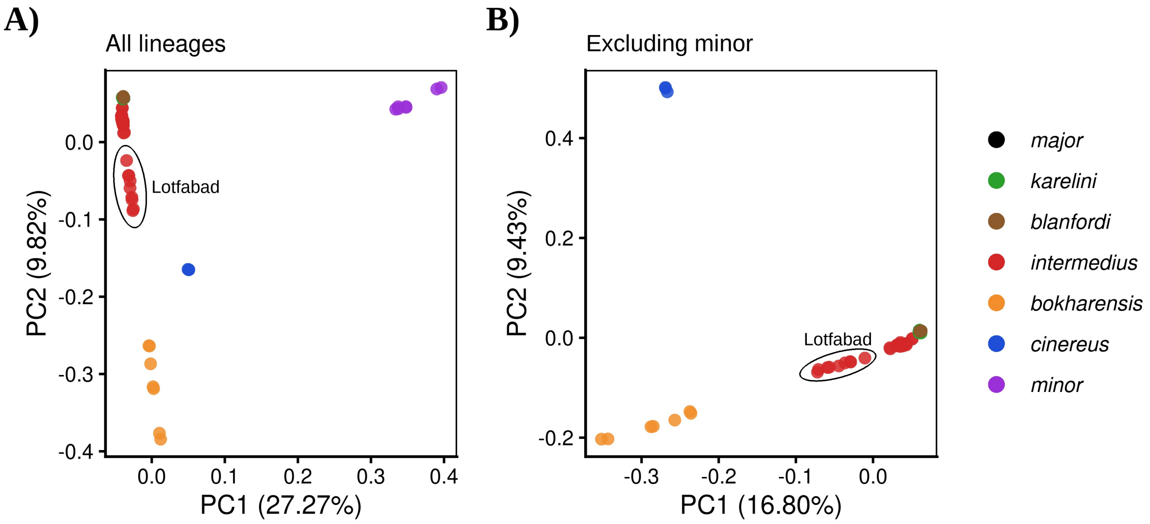


**Figure S4: Principal component analysis of the *Parus major* complex.** PC1 versus PC2 for **(A)** all lineages (87 individuals) and **(B)** all lineages excluding *minor* (80 individuals), based on biallelic SNPs thinned to $\geq$1 kb spacing; individuals with >10% missing genotypes were excluded (see Methods). Ellipses enclose admixed individuals from Lotfabad (Fig. 1).

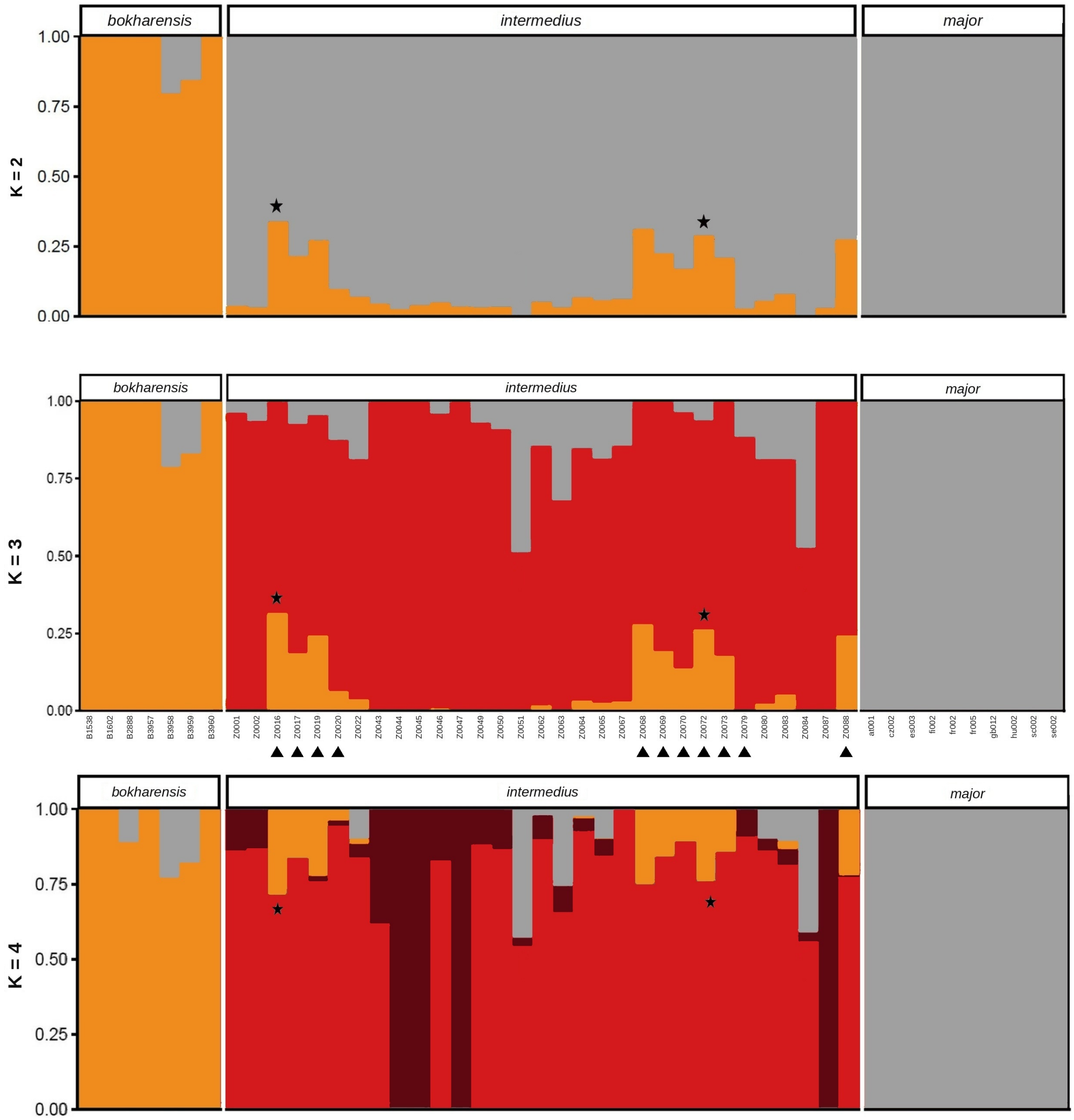


**Figure S5: Individual ancestry proportions inferred using ADMIXTURE at $K = 2$–$4$.** Each vertical bar represents one individual, grouped by population; sample IDs (shown below the $K = 3$ panel) apply to all panels. Orange, *bokharensis*; grey, *major*; light red, main *intermedius* component ($K = 3$ and $K = 4$); dark red, second *intermedius* component ($K = 4$). Triangles below the sample IDs mark *intermedius* individuals sampled at Lotfabad, in the putative contact zone between the *major* and *bokharensis* lineages; stars mark the two Lotfabad individuals carrying *bokharensis* mitochondrial haplotypes (Z0016, Z0072). Cross-validation errors for each value of $K$ are reported in Table S4.

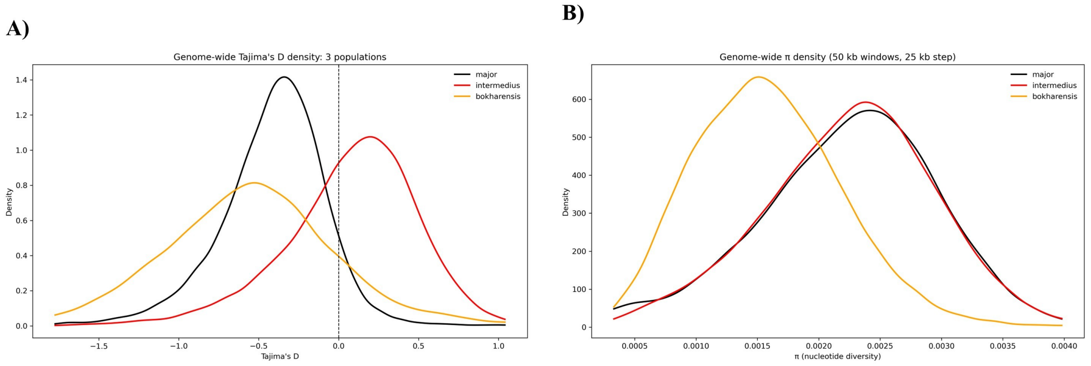


**Figure S6: Genome-wide distributions of nucleotide diversity and Tajima's $D$ across three focal populations.** (A) Density curves of Tajima's $D$ calculated in 50 kb sliding windows (25 kb step) across the genome. (B) Density curves of nucleotide diversity ($\pi$) calculated across the genome. Populations are shown as *intermedius* (red), *bokharensis* (orange), and *major* (black). The dashed line in (A) indicates neutrality (Tajima's $D = 0$).

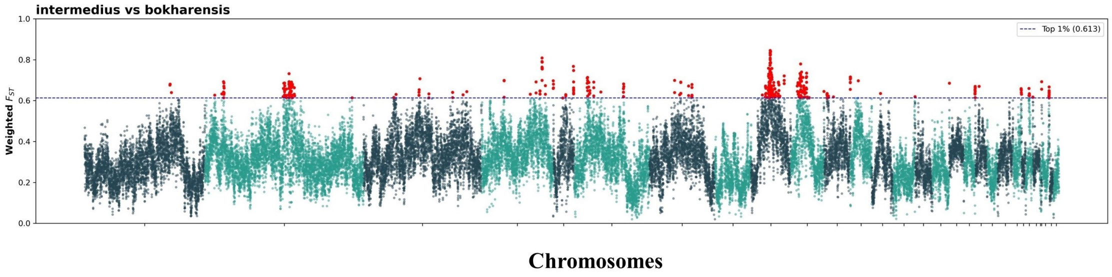


**Figure S7: Genome-wide distribution of $F_{\mathrm{ST}}$ between *P. bokharensis* and *P. m. intermedius*.** Points represent sliding windows (50 kb window size, 25 kb step size) across the genome, with outlier windows highlighted in red; the dashed line indicates the top 1% empirical threshold.

### Supplementary Tables

**Table S1:** Sample metadata, whole-genome sequencing coverage and dataset membership for all individuals analysed in this study. Check marks indicate whether each individual was included in nuclear analyses ($n = 112$) and in mitogenome analyses ($n = 99$; Fig. S2).

| Taxon | Sample ID | Locality | Lat. | Long. | Sex | Coverage | Nuclear | Mitogenome |
|---|---|---|---|---|---|---|---|---|
| *P. m. intermedius* | Z0064 | Dargaz | 37.39 | 58.87 | F | 21.599 | ✓ | ✓ |
| | Z0020 | Lotfabad | 37.42 | 59.52 | M | 21.2814 | ✓ | ✓ |
| | Z0080 | Dargaz | 37.39 | 58.87 | F | 17.1873 | ✓ | ✓ |
| | Z0063 | Dargaz | 37.39 | 58.87 | F | 16.4539 | ✓ | ✓ |
| | Z0084 | Bojnurd | 37.47 | 57.33 | M | 19.7359 | ✓ | ✓ |
| | Z0022 | Dargaz | 37.39 | 58.87 | F | 19.265 | ✓ | ✓ |
| | Z0001 | Kardeh dam | 36.67 | 59.53 | F | 21.4507 | ✓ | ✓ |
| | Z0072 | Lotfabad | 37.42 | 59.52 | F | 22.2876 | ✓ | ✓ |
| | Z0045 | Ortokand | 37.65 | 59.29 | M | 18.3558 | ✓ | ✓ |
| | Z0070 | Lotfabad | 37.42 | 59.52 | M | 21.0484 | ✓ | ✓ |
| | Z0049 | Mashhad | 36.30 | 59.61 | M | 17.5149 | ✓ | ✓ |
| | Z0062 | Dargaz | 37.39 | 58.87 | M | 17.4717 | ✓ | ✓ |
| | Z0073 | Lotfabad | 37.42 | 59.52 | M | 20.6154 | ✓ | ✓ |
| | Z0065 | Dargaz | 37.39 | 58.87 | M | 16.9305 | ✓ | ✓ |
| | Z0043 | Ortokand | 37.65 | 59.29 | M | 17.9301 | ✓ | ✓ |
| | Z0083 | Dargaz | 37.39 | 58.87 | F | 17.4721 | ✓ | ✓ |
| | Z0051 | Bojnurd | 37.47 | 57.33 | M | 19.2622 | ✓ | ✓ |
| | Z0088 | Lotfabad | 37.42 | 59.52 | M | 17.8272 | ✓ | ✓ |
| | Z0050 | Moghan | 36.13 | 59.37 | M | 20.3758 | ✓ | ✓ |
| | Z0046 | Ortokand | 37.65 | 59.29 | M | 47.7351 | ✓ | ✓ |
| | Z0067 | Dargaz | 37.39 | 58.87 | M | 18.8132 | ✓ | ✓ |
| | Z0016 | Lotfabad | 37.42 | 59.52 | M | 17.6469 | ✓ | ✓ |
| | Z0087 | Ortokand | 37.65 | 59.29 | F | 19.6493 | ✓ | ✓ |
| | Z0079 | Lotfabad | 37.42 | 59.52 | M | 19.3017 | ✓ | ✓ |
| | Z0019 | Lotfabad | 37.42 | 59.52 | F | 24.2285 | ✓ | ✓ |
| | Z0002 | Kardeh dam | 36.67 | 59.53 | M | 23.4098 | ✓ | ✓ |
| | Z0069 | Lotfabad | 37.42 | 59.52 | M | 16.7805 | ✓ | ✓ |
| | Z0047 | Ortokand | 37.65 | 59.29 | M | 19.2281 | ✓ | ✓ |
| | Z0044 | Ortokand | 37.65 | 59.29 | F | 17.1366 | ✓ | ✓ |
| | Z0068 | Lotfabad | 37.42 | 59.52 | M | 23.3577 | ✓ | ✓ |
| | Z0017 | Lotfabad | 37.42 | 59.52 | M | 22.1196 | ✓ | ✓ |
| *P. m. karelini* | Z0029 | Gorgan | 36.78 | 54.46 | M | 20.023 | ✓ | ✓ |
| | Z0009 | Ardebil | 37.33 | 48.40 | F | 22.8296 | ✓ | ✓ |
| | Z0024 | Gorgan | 36.78 | 54.46 | M | 45.438 | ✓ | ✓ |
| | Z0042 | Noor | 36.57 | 52.05 | M | 46.0332 | ✓ | ✓ |
| | Z0037 | Noor | 36.57 | 52.05 | M | 46.5165 | ✓ | ✓ |
| | Z0007 | Ardebil | 37.33 | 48.40 | M | 46.734 | ✓ | ✓ |
| | Z0026 | Gorgan | 36.78 | 54.46 | F | 49.7561 | ✓ | ✓ |
| | Z0010 | Ardebil | 37.33 | 48.40 | M | 20.7957 | ✓ | ✓ |
| | Z0041 | Noor | 36.57 | 52.05 | M | 22.3487 | ✓ | ✓ |
| | Z0006 | Zanjan | 36.55 | 47.62 | M | 17.5468 | ✓ | ✓ |
| | Z0086 | Gorgan | 36.78 | 54.46 | F | 43.26 | ✓ | ✓ |
| | Z0060 | Orumiyeh | 37.55 | 45.12 | F | 17.1274 | ✓ | ✓ |
| | Z0025 | Gorgan | 36.78 | 54.46 | M | 49.7911 | ✓ | ✓ |
| | Z0040 | Noor | 36.57 | 52.05 | M | 46.6942 | ✓ | ✓ |

| Taxon | Sample ID | Locality | Latitude | Longitude | Sex | Coverage | Nuclear | Mitogenome |
|---|---|---|---|---|---|---|---|---|
| | Z0005 | Zanjan | 36.55 | 47.62 | M | 45.9884 | ✓ | – |
| | Z0004 | Zanjan | 36.55 | 47.62 | M | 20.8865 | ✓ | ✓ |
| | Z0027 | Gorgan | 36.78 | 54.46 | M | 55.6598 | ✓ | ✓ |
| | Z0023 | Gorgan | 36.78 | 54.46 | F | 22.3696 | ✓ | ✓ |
| | Z0028 | Gorgan | 36.78 | 54.46 | F | 40.6163 | ✓ | ✓ |
| | Z0031 | Gorgan | 36.78 | 54.46 | F | 48.5936 | ✓ | ✓ |
| | Z0081 | Orumiyeh | 37.55 | 45.12 | F | 17.4457 | ✓ | ✓ |
| | Z0038 | Noor | 36.57 | 52.05 | F | 44.4248 | ✓ | ✓ |
| | Z0039 | Noor | 36.57 | 52.05 | F | 22.3463 | ✓ | ✓ |
| | Z0033 | Noor | 36.45 | 51.79 | M | 45.5955 | – | ✓ |
| | Z0034 | Noor | 36.45 | 51.79 | M | 53.0923 | – | ✓ |
| | Z0035 | Noor | 36.45 | 51.79 | F | 18.1401 | – | ✓ |
| *P. m. blanfordi* | Z0011 | Yasuj | 30.67 | 51.59 | F | 18.4913 | ✓ | ✓ |
| | Z0076 | Yasuj | 30.67 | 51.59 | M | 17.1616 | ✓ | ✓ |
| | Z0056 | Khuzestan | 31.32 | 48.67 | M | 17.8242 | ✓ | ✓ |
| | Z0012 | Yasuj | 30.67 | 51.59 | M | 46.0528 | ✓ | ✓ |
| | Z0014 | Yasuj | 30.67 | 51.59 | F | 48.9821 | ✓ | ✓ |
| | Z0057 | Khuzestan | 31.32 | 48.67 | F | 19.3085 | ✓ | ✓ |
| | Z0074 | Shiraz | 29.59 | 52.58 | M | 21.418 | ✓ | ✓ |
| | Z0055 | Khuzestan | 31.32 | 48.67 | F | 48.1251 | ✓ | ✓ |
| | Z0075 | Tabriz | 38.08 | 46.29 | M | 18.9205 | ✓ | ✓ |
| | Z0077 | Yasuj | 31.24 | 51.18 | F | 18.2254 | – | ✓ |
| *P. major* | SRR5423293 | Oxford, United Kingdom | 51.75 | -1.26 | M | 53.6982 | ✓ | ✓ |
| | SRR5423294 | Corsica, France | 42.04 | 9.01 | M | 58.6243 | ✓ | ✓ |
| | SRR5423295 | La Rouvière, France | 43.94 | 3.73 | M | 50.1128 | ✓ | ✓ |
| | SRR5423296 | Velký Kosíř, Czech Republic | 49.55 | 17.04 | M | 55.0028 | ✓ | ✓ |
| | SRR5423297 | Oulu, Finland | 65.01 | 25.47 | M | 64.4316 | ✓ | ✓ |
| | SRR5423298 | Loch Lomond, Scotland | 56.08 | -4.58 | M | 55.1742 | ✓ | ✓ |
| | SRR5423299 | Gotland, Sweden | 57.47 | 18.49 | M | 55.5337 | ✓ | ✓ |
| | SRR5423300 | Pilis Mountains, Hungary | 47.69 | 18.89 | M | 61.0345 | ✓ | ✓ |
| | SRR5423301 | Austria | 48.06 | 15.91 | M | 53.2034 | ✓ | ✓ |
| | SRR5423302 | Sierra de Mariola, Spain | 38.76 | -0.53 | M | 52.7807 | ✓ | ✓ |
| | SRR1793425 | Eerguna, Inner Mongolia, China | 51.97 | 120.55 | F | 4.8762 | – | ✓ |
| | SRR1793427 | Eerguna, Inner Mongolia, China | 51.97 | 120.55 | M | 5.2181 | – | ✓ |
| | SRR1793428 | Eerguna, Inner Mongolia, China | 51.97 | 120.55 | M | 5.0058 | – | ✓ |
| | SRR1793429 | Eerguna, Inner Mongolia, China | 51.97 | 120.55 | F | 3.9286 | – | ✓ |
| | SRR1793430 | Teshig, Mongolia | 49.24 | 102.27 | M | 4.6511 | – | ✓ |
| | SRR1793431 | Teshig, Mongolia | 49.24 | 102.27 | F | 5.2329 | – | ✓ |
| | SRR1793432 | Tushig, Mongolia | 50.52 | 105.00 | M | 4.3378 | – | ✓ |
| | SRR1793433 | Tushig, Mongolia | 50.52 | 105.00 | M | 4.1743 | – | ✓ |
| | SRR1793434 | Tushig, Mongolia | 50.52 | 105.00 | F | 4.1959 | – | ✓ |
| *P. bokharensis* | B1538 | Yarodar, Kyrgyzstan | 41.00 | 75.00 | M | 41.804 | ✓ | ✓ |

| Taxon | Sample ID | Locality | Latitude | Longitude | Sex | Coverage | Nuclear | Mitogenome |
|---|---|---|---|---|---|---|---|---|
| | B1602 | Ili River near Kapchagay, Kazakhstan | 43.87 | 77.05 | M | 32.3945 | ✓ | ✓ |
| | B2888 | Chokpak, Kazakhstan | 42.62 | 70.58 | M | 35.1339 | ✓ | ✓ |
| | B3957 | Chokpak, Kazakhstan | 42.62 | 70.58 | F | 37.1073 | ✓ | ✓ |
| | B3958 | Chokpak, Kazakhstan | 42.62 | 70.58 | F | 40.2623 | ✓ | ✓ |
| | B3959 | Chokpak, Kazakhstan | 42.62 | 70.58 | M | 40.0375 | ✓ | ✓ |
| | B3960 | Chokpak, Kazakhstan | 42.62 | 70.58 | F | 142.197 | ✓ | – |
| | B3961 | Chokpak, Kazakhstan | 42.62 | 70.58 | M | 36.21 | – | ✓ |
| *P. minor* | SRR1793405 | Yanbian, Sichuan | 26.47 | 101.60 | M | 4.53487 | ✓ | – |
| | SRR1793406 | Yanbian, Sichuan | 24.98 | 98.74 | M | 4.70339 | ✓ | – |
| | SRR1793407 | Gaoligong, Yunnan | 24.98 | 98.74 | M | 5.49632 | ✓ | – |
| | SRR1793415 | Gaoligong, Yunnan | 24.98 | 98.74 | F | 5.55623 | ✓ | – |
| | SRR1793417 | Gaoligong, Yunnan | 24.98 | 98.74 | M | 5.88094 | ✓ | – |
| | SRR1793419 | Gaoligong, Yunnan | 24.98 | 98.74 | F | 5.2391 | ✓ | – |
| | SRR1793420 | Gaoligong, Yunnan | 24.98 | 98.74 | M | 4.69342 | ✓ | – |
| | SRR1793421 | Lijiang, Yunnan | 27.10 | 100.07 | F | 4.4292 | ✓ | – |
| | SRR1793422 | Lijiang, Yunnan | 27.10 | 100.07 | M | 4.95103 | ✓ | – |
| | SRR1793423 | Lijiang, Yunnan | 27.10 | 100.07 | F | 4.4587 | ✓ | – |
| | SRR1793424 | Lijiang, Yunnan | 27.10 | 100.07 | M | 4.25471 | ✓ | – |
| | SRR1793436 | Shennongjia, Hubei | 31.88 | 110.92 | F | 4.91001 | ✓ | – |
| | SRR1793437 | Shennongjia, Hubei | 31.88 | 110.92 | M | 4.97231 | ✓ | – |
| | SRR1793438 | Jixi, Anhui | 30.20 | 115.54 | M | 5.24855 | ✓ | – |
| | SRR1793439 | Panjin, Liaoning | 40.65 | 121.46 | M | 5.1524 | ✓ | – |
| | SRR1793440 | Panjin, Liaoning | 40.65 | 121.46 | M | 4.68994 | ✓ | ✓ |
| | SRR1793441 | Panjin, Liaoning | 40.65 | 121.46 | M | 4.89532 | ✓ | – |
| | SRR1793442 | Kuandian, Liaoning | 40.98 | 125.43 | M | 4.07455 | ✓ | – |
| | SRR1793443 | Kuandian, Liaoning | 40.98 | 125.43 | M | 4.58637 | ✓ | ✓ |
| | SRR1793444 | Sunan, Gansu | 37.84 | 102.01 | F | 5.07372 | ✓ | – |
| | SRR1793445 | Sunan, Gansu | 37.84 | 102.01 | M | 5.08291 | ✓ | – |
| | SRR1793447 | Yongdeng, Gansu | 36.68 | 102.73 | F | 4.94089 | ✓ | – |
| | SRR765717 | Lijiang, Yunnan | 27.10 | 100.07 | F | 12.9284 | ✓ | – |
| | SRR9656355 | Foping, Shaanxi | 33.52 | 107.98 | M | 16.456 | ✓ | ✓ |
| | SRR9656356 | Libo, Guizhou | 25.26 | 108.05 | F | 12.1379 | ✓ | – |
| | SRR9656357 | Chaoan, Guangdong | 23.71 | 116.54 | M | 13.6619 | ✓ | ✓ |
| | SRR9656367 | Baishan, Jilin | 41.86 | 126.52 | M | 13.1701 | ✓ | – |
| | SRR9656368 | Huairou, Beijing | 40.37 | 116.67 | F | 16.261 | ✓ | – |
| | SRR9656369 | Mangkang, Tibet | 29.27 | 98.69 | M | 15.2459 | ✓ | – |
| *P. cinereus* | C2761 | Nepal | 27.19 | 87.10 | M | 35.867 | ✓ | ✓ |
| | C2762 | Nepal | 27.19 | 87.10 | F | 38.4023 | ✓ | ✓ |
| | C4242 | Nepal | 27.99 | 85.20 | F | 39.776 | ✓ | ✓ |

**Table S2: Sample sizes and inclusion rules for each genomic analysis.** Range-wide nuclear data set, 112 individuals; regional data set, 48 individuals (*major*, *intermedius*, *bokharensis*); mitogenome data set, 99 individuals; 126 individuals in total.

| Analysis | $n$ | Lineages | Inclusion rule |
|---|---|---|---|
| **Phylogeny and genome-wide introgression** | | | |
| Mitogenome tree and network | 99 | all | assembled mitogenome |
| Nuclear species tree (ASTRAL) | 12 | all | 1–2 high-coverage males per lineage |
| $D$, $f_4$-ratio, $f$-branch (Dsuite) | 12 | all | as species tree |
| **Demographic history** | | | |
| MSMC2 | 2 | *major*, *bokharensis* | one high-coverage (>20×) male per lineage |
| **Population structure and admixture** | | | |
| PCA, all lineages | 87 | all | range-wide; ≤10% missing data |
| PCA, excluding *minor* | 80 | all but *minor* | as above |
| PCA, ADMIXTURE, hybrid index | 48 | focal | regional, all |
| **Differentiation and diversity** | | | |
| Genome-wide $F_{\mathrm{ST}}$, $\pi$, Tajima's $D$ | 112 | all | range-wide, all |
| Sliding-window $F_{\mathrm{ST}}$, $d_{XY}$, $\pi$, Tajima's $D$ | 43 | focal | regional, reference subset[a] |
| Coding diversity ($\pi_0/\pi_4$; ANGSD) | 73 | *major* lineage[b], *intermedius* | all *intermedius* (31) and *major*-lineage (42) individuals |
| **Localized introgression (chromosome 24)** | | | |
| $f_{dM}$, TWISST, DIEM | 44 | focal, *minor* | reference subset[a] + 1 *minor* outgroup |

Focal: *major*, *intermedius* and *bokharensis*. [a] Excluding two *bokharensis* individuals with *major* ancestry (16–20%) and two *intermedius* individuals without detectable *bokharensis* ancestry (<1%) at $K = 2$ (ADMIXTURE), leaving nine *major*, 29 *intermedius* and five *bokharensis* individuals. [b] European *major* ($n = 10$), *karelini* ($n = 23$) and *blanfordi* ($n = 9$).

**Table S3: Number of loci across the genome using different filtering settings for 10 kb non-overlapping windows.** Filtering parameters include minimum read depth (DP), minimum percentage of the window covered by data (PW), and missing data per site (MD).

| Filtering setting | Number of total trees |
|---|---|
| 1) DP = 1, PW = 50%, MD = 15% | 91100 |
| 2) DP = 5, PW = 50%, MD = 15% | 90549 |
| 3) DP = 1, PW = 50%, MD = 5% | 90920 |
| 4) DP = 1, PW = 80%, MD = 10% | 72988 |

**Table S4: ADMIXTURE cross-validation results.** Cross-validation errors for ADMIXTURE analyses conducted at $K = 2$–$4$ ancestral clusters.

| $K$ | Cross-validation error |
|---|---|
| 2 | 0.582 |
| 3 | 0.606 |
| 4 | 0.643 |

**Table S5:** Genes located within the highly differentiated region on chromosome 24 surrounding the *BCO2* locus.

| Gene | Gene coordinates (chr24) | Number of SNPs ($F_{ST} > 0.9$) | Biological function |
|---|---|---|---|
| *BCO2* | 1480773–1495350 | 8 | Carotenoid metabolism enzyme controlling pigment degradation (GeneCards) |
| *TEX12* | 1497855–1502348 | 9 | Meiosis-related protein important in gamete formation |
| *LOC107214402* | 1502582–1506640 | 9 | Predicted protein (uncharacterized) |
| *SDHD* | 1507725–1511065 | 1 | Mitochondrial respiratory chain protein |
| *NKAPD1* | 1511670–1515119 | 1 | Nuclear protein involved in transcription regulation (GeneCards) |
| *DLAT* | 1516079–1524391 | 9 | Enzyme of the pyruvate dehydrogenase complex involved in energy metabolism (GeneCards) |
| *DIXDC1* | 1525317–1551085 | 2 | Wnt signaling pathway protein |

**Table S6:** Comparison of the two highly differentiated *BCO2* 5′ UTR variants (NC_031792.1:1495013 G/A and NC_031792.1:1494978 A/G) across selected parid species. Homologous sequence was recovered only for closely related taxa. The complete haplotype associated with *P. bokharensis* was observed only within *P. bokharensis* and *P. major intermedius*, whereas other parid (sub)species carried either the major haplotype or mixed combinations of the two alleles.

| Taxon | 1495013 | 1494978 | Haplotype |
|---|---|---|---|
| *Parus major* | G | A | major |
| *Parus minor* | G | A | major |
| *Parus major intermedius* | A | G | bokharensis-like |
| *Parus bokharensis* | A | G | bokharensis-like |
| *Parus cinereus* | G/A | A | major/Mixed |
| *Poecile atricapillus* | G | G | Mixed |
| *Cyanistes caeruleus* | G | G | Mixed |
| *Pseudopodoces humilis* | NA | A | major-like (partial) |
